# A comparison of adiponectin-deficient mice reveals the fundamental role of intracellular adiponectin

**DOI:** 10.1101/2025.07.12.664095

**Authors:** Toshiharu Onodera, Dae-Seok Kim, May-Yun Wang, Megan Virostek, Shiuhwei Chen, Zachary A Kipp, Evelyn A Bates, Yan Li, Clair Crewe, Chao Li, Xue-Nan Sun, Qianbin Zhang, Mengle Shao, Jihoon Shin, Christine M Kusminski, Ruth Gordillo, Rana K Gupta, Terry D Hinds, Iichiro Shimomura, Philipp E Scherer

**Affiliations:** Touchstone Diabetes Center, The University of Texas Southwestern Medical Center, Dallas, TX, USA; Department of Metabolic Medicine, Osaka University Graduate School of Medicine, Suita, Osaka, Japan; Department of Adipose Management, Osaka University Graduate School of Medicine, Osaka, Japan; Department of Pharmacology and Nutritional Sciences, University of Kentucky College of Medicine, Lexington, Kentucky, USA; Drug & Disease Discovery D3 Research Center, Department of Pharmacology and Nutritional Sciences, University of Kentucky College of Medicine, Lexington, Kentucky, USA; Markey Cancer Center, University of Kentucky, Lexington, Kentucky, USA; Barnstable Brown Diabetes Center, University of Kentucky College of Medicine, Lexington, Kentucky, USA; Department of Cell Biology and Physiology, Washington University School of Medicine, St. Louis, MO, USA and Department of Internal Medicine, Division of Endocrinology, Metabolism and Lipid Research, Washington University School of Medicine, St. Louis, MO, USA; CAS Key Laboratory of Molecular Virology and Immunology, The Center for Microbes, Development and Health, Chinese Academy of Sciences, Shanghai, China; Division of Endocrinology, Diabetes and Metabolism, Beth Israel Deaconess Medical Center and Harvard Medical School, Boston, MA, USA; Division of Endocrinology, Department of Medicine, Duke Molecular Physiology Institute, Duke University, Durham, NC, USA

**Keywords:** adiponectin, adipose tissue

## Abstract

Adiponectin is an important adipokine with insulin-sensitizing, anti-inflammatory, and anti-fibrotic properties. The physiological roles of adiponectin have been studied using global adiponectin knockout (KO) mice. However, the reported phenotypes of adiponectin KO mice vary based on the mouse lines generated by different strategies and investigators.

We performed a head-to-head comparison of the adiponectin KO mice that were generated in Dallas, Houston and Osaka. RNAseq revealed that the expression of the bioactive domain of adiponectin – the globular domain - was preserved in the Houston and Osaka KO mice. A complete adiponectin KO model, such as the Dallas KO mouse, exhibits a lower body weight, the highest adipocyte mitochondrial function and displays a susceptibility to DNA damage-mediated lung fibrosis. The reconstitution of globular adiponectin into the Dallas KO mice prompted an increase in body weight and a partial recapitulation of the Osaka KO model transcriptome signature. The intracellular globular adiponectin form is important, as we found that globular adiponectin enhances PPARγ activity by modulating the coregulators interacting with PPARγ.

Overall, the residual expression of globular adiponectin regulates adipose tissue metabolism by altering PPARγ activity, highlighting an important novel role of intracellular adiponectin.

## Introduction

Adiponectin is a protein predominantly produced by adipocytes that exerts pleiotropic effects on the heart, liver, kidney, and pancreas, as well as endothelial and immune cells ^1–5^. Adiponectin is a secretory protein interacting with adiponectin receptors (AdipoRs), activates the receptor-inherent or receptor-associated ceramidase activity in the AdipoRs and targets downstream molecules, such as AMPK. AMP-activated protein kinase (AMPK) was originally identified as a signal transducer of AdipoR1 in muscle^6^. Independent of AMPK activity, AdipoR1 and AdipoR2 display ceramidase activity and / or associated with a ceramidase and thereby contribute to the reduction of cytotoxic ceramide lipid species^7,8^.

Adiponectin function has been explored in many different settings utilizing recombinant adiponectin protein ^9^, overexpression of adiponectin by viruses ^10^, adiponectin receptor agonists ^11^, as well as genetic mouse models ^12,13^. Importantly, the studies conducted in adiponectin-deficient mice contributed to a better understanding of the physiological roles of adiponectin. The generation of whole-body adiponectin deficient mice was attempted in at least four places, including in institutions located in Dallas, Osaka, Tokyo, and Houston ^12,14–16^. The first report on the Houston adiponectin KO mice exhibited relatively lower insulin levels during oral glucose tolerance test (OGTT) and higher fatty acid oxidative capacity ^14^. On the contrary, the Osaka adiponectin KO mice showed impaired glucose tolerance and lipid metabolism ^15^. The glucose tolerance in the Dallas adiponectin KO mice varied by context. Hepatic insulin sensitivity is disrupted, but one report shows improved serum glucose and insulin levels in OGTTs ^12,17^. In line with this, the Tokyo adiponectin KO mice originally showed moderate glucose intolerance and decreased insulin signaling but improved hepatic and muscle steatosis ^16,18^. A follow-up study by the Tokyo group demonstrated that adiponectin KO mice displayed reduced body weight gain, reduced leptin, and higher energy expenditure under high fat diet (HFD) conditions ^19^.

In addition to the parameters describing glucose and lipid metabolism, the phenotypic manifestations in various organs also vary greatly among the different kinds of adiponectin KO mice. This occurs despite the fact that adiponectin is produced predominantly in adipocytes and the pool of *circulating* adiponectin is lost equally in all adiponectin KO mice. Adiponectin is protective from caspase-mediated podocyte ablation as shown in the Dallas adiponectin KO mice ^4^, while adiponectin reportedly becomes deleterious to the development of renal fibrosis ^20^ due to proinflammatory effects of adiponectin in the Houston adiponectin KO mice ^21^. Additionally, the deficiency of adiponectin affects the thermogenic phenotype of subcutaneous adipose tissue in various ways. Adiponectin suppresses the browning of subcutaneous adipose tissue in Dallas and in Tokyo adiponectin KO mice ^19,22^. However, adiponectin enhances thermogenesis in the Houston adiponectin KO mice by mediating M2 macrophage accumulation ^23^.

This raises a number of questions. Where do these phenotypic discrepancies among adiponectin KO mice stem from? The answer to this question has the potential to shed significant light on the role of adiponectin within the cell that produces it.

Mouse adiponectin protein is composed of 247 amino acids with 4 domains: an N-terminal signal sequence, a variable region, a collagenous domain of 22 G-X-Y repeats, and a C-terminal globular trimerization domain (the “globular domain”) ^9^. At the genomic level, the mouse adiponectin locus consists of 3 exons. Exon 1 contains the 5’-UTR. Exon 2 contains the start codon and the signal sequence that is required for export into the extracellular milieu. It also encodes the variable domain and a portion of the collagenous domain. Exon 3 includes the rest of the collagenous domain as well as the globular domain, which is essential for the bioactivity of adiponectin ^24^. The basic strategy of the elimination of adiponectin in both the Houston and the Osaka adiponectin KO mouse is the disruption of Exon 2, which contains the translation initiation codon ^14,15^. This is, by all accounts, a sensible way to disrupt the gene, particularly for a disruption affecting a secretory protein. However, exon1, and the most functional part of adiponectin, exon3, remain intact in these models. In contrast, the deletion of the whole adiponectin gene is accomplished in the Dallas model by replacing all three exons and the 5’-untranslated region of the adiponectin gene with a neomycin resistance cassette ^12^.

We hypothesized that the different knockout strategies are responsible for the differences of phenotypes observed in the three adiponectin KO mice. Since the Osaka and Houston adiponectin KO mice lack only exon 2, the unexpected translation of exon 3 might result in the production of globular adiponectin. We appreciate that the deletion of exon 2 eliminates the ability of adiponectin to be secreted because exon 2 includes the signal sequence that guides the proteins through the secretory pathway to the extracellular destinations. However, the residual expression of exon 3 of adiponectin may preserve the biological activity of adiponectin since exon 3 encodes the globular domain that gives rise to a bioactive form of adiponectin ^24,25^. Therefore, we wanted to test specifically whether the intracellular accumulation of the exon3-encoded portion of adiponectin can explain the phenotypic differences specifically in Osaka and Houston adiponectin KO mice. This is also important since we increasingly appreciate the role of local adiponectin expression in the kidney, the hepatic stellate cell, and the cardiomyocyte.

To settle the discrepancies of the phenotypes derived from the differential adiponectin KO approaches, we took advantage of adiponectin deficient mice generated in Dallas, Houston and Osaka by breeding them into the same genetic background and performing simultaneous experiments in all three models, performed under identical conditions. We performed head-to-head comparisons of adiponectin-deficient mice to confirm the phenotypes that have been reported previously and aimed to expand the repertoire of adiponectin-associated phenotypes by designing new experiments allowing us to cross-compare. We also generated and analyzed a mouse overexpressing intracellular globular adiponectin that corresponds to the residual portion of adiponectin in the Houston and Osaka adiponectin KO mice. Through these analyses, we describe the potential mechanistic reasons for the phenotypic differences of adiponectin in these KO mouse lines and clarify for the first time the differential role of intracellular and extracellular adiponectin.

## Research Design and Methods

### Mice

The transgenic mouse strain *Dallas adiponectin deficient mice* were generated and described by our laboratory previously ^12^. The *Houston adiponectin deficient mice* were purchased from Jackson laboratory ^14^ and have been used widely in the literature. The *Osaka adiponectin deficient mice* were kindly provided by Dr. Iichiro Shimomura ^15^. All mice were bred to a C57BL/6 genetic background. Mice were fed on regular (LabDiet #5058) or high-fat diet (60%, Research #D12492). Mice were maintained in 12-h dark/light cycles, with *ad libitum* access to diet and water. All protocols for mouse experiments and euthanasia were reviewed and approved by the Institutional Animal Care and Use Committee of the University of Texas Southwestern Medical Center, animal protocol number 2015-101207G.

### Genotyping PCR

Approximately 3 mm of mouse tail tip was incubated in 80 μL 50 mM NaOH at 95 °C for 1.5 h. 8 μL 1 M Tris–HCl (pH 8.0) was added for neutralization. After vortexing and a short spin down, 0.5–1 μL of supernatant was used as <u>PCR</u> template. Primer sequences for genotyping PCR are listed in <u>Table S1</u>. The PCR program was: 95 °C for 5 min, followed by 35 cycles of 95 °C for 15 s, 62 °C for 30 s, and 72 °C for 30 s, and ended with 72 °C for 3 min.

### Immunohistochemistry

Mice were euthanized by cervical dislocation following isoflurane anesthesia. Tissues were immediately collected and fixed in 10% buffered formalin for 24 hours. Afterwards, tissues were rinsed with 50% ethanol and embedded in paraffin blocks and sliced into 5 μm sections. H&E staining (#ab245880, Abcam) and Trichrome staining (#ab150686, Abcam) were performed according to the manufacturer’s instructions. H&E stained and Trichrome stained images were taken by Keyence BZ-X710 microscope (Keyence). Adipocyte size was measured with the Keyence BZ-X Analyzer software ^26^.

### Assay of metabolites

Tail blood was collected with a Microvette CB300Z (SARSTEDT #16.440.100) for serum and a Microvette CB300 K2E (SARSTEDT #16.444.100) was used for plasma collection. Glucose was measured with a glucose meter or with colorimetric assays with PGO enzymes (Sigma #P7119) plus o-dianisidine (Sigma #F5803). Insulin was measured with an ELISA kit (Crystal Chem #90080). Urine citrate was measured by enzymatic citrate lyase ^27^. Urine NH_4_ was measured with Infinity Ammonia Reagent (Beckman Coulter #OSR61154).

### qPCR

Total RNA was extracted with an RNeasy Mini kit (Qiagen #74106) for pancreas or Trizol (Invitrogen #15596018) for kidney, liver, adipose tissue and other organ. cDNA was synthesized with iScript cDNA Synthesis Kit (Bio-Rad #170-8891). Quantitative real-time PCR (qPCR) was performed with the Powerup SYBR Green PCR master mix (Applied Biosystems # A25742) on Quantistudio 6 Flex Real-Time PCR System (Applied Biosystems # 4485694). Primer sequences for qPCR are listed in Table S2.

### Western blotting

Protein was extracted from adipose tissue by homogenization in PBS supplemented with 1 mM EDTA, 20 mM NaF, 2 mM Na_3_VO_4_, and protease inhibitor cocktail (539131, EMD Millipore). 5× RIPA buffer was added to the homogenate for a final concentration of 10 mM Tris-HCl, 2 mM EDTA, 0.3% NP40, 0.3% deoxycholate, 0.1% SDS, and 140 mM NaCl, pH 7.4. The sample was centrifuged at 10,000 *g* for 5 minutes. 20–50 μg/lane of supernatant protein was separated by SDS-PAGE (NP0335BOX, Thermo Fisher) and transferred to a nitrocellulose membrane. The blots were then incubated overnight at 4°C with rabbit anti-mouse polyclonal adiponectin antibodies (produced in-house), ADP-ribose binding reagent (MABE1031, EMD Millipore), PARP1 antibody (39559, Active Motif), FLAG antibody (F1804, Sigma or 14793, Cell signaling), PPARγ antibody (sc-7273, Santa Cruz Biotechnology), GAPDH antibody (5174, Cell Signaling) in a 1% BSA TBST-blocking solution, respectively. Primary antibodies were detected using secondary antibodies labeled with infrared dyes emitting at 700 nm or 800 nm (1:5000, Li-Cor Bioscience 925-68073 and 926-32213, respectively). The Odyssey Infrared Imager was used to visualize Western blots.

### Immunoprecipitation

Adipose tissue lysates were prepared as described in the <u>Western blot</u> section. The anti-FLAG antibody (14793, Cell Signaling Technology) was bound to protein A and G magnetic beads (BioRad, cat#: 161-4013 and 161-4023) in TBST (50mM Tris-HCl, 0.1 mM EDTA, 150 mM NaCl and 0.5% Tween-20, pH 7.5). Anti-FLAG beads (M8823, Sigma) were used for the precipitation of FLAG protein. 100 μg protein lysate was incubated with the anti-FLAG beads overnight at 4°C. Beads were washed 3 times with TBST and eluted in 0.1M glycine buffer at pH2.5 for 10 min at room temperature. Proteins were then separated by SDS-PAGE. Secondary antibodies and detection were described in the Western blot section.

### Systemic tests

For oral glucose tolerance tests (OGTTs), mice were fasted for 4–6 h and subjected to an oral gavage of dextrose (2.5 mg/g body weight). Tail blood was collected at 0, 15, 30, 60, and 120 min and prepared for serum and assayed for glucose and insulin. For insulin tolerance test (ITT), insulin (1.5 U/kg Humulin R; Eli Lilly, Indianapolis, IN, USA) was administered under fed condition. Serum glucose level was measured at the 0, 15, 30, 60, and 120 minutes time point. Triglyceride tolerance test (TGTT) was initiated by oral gavage of 20% Intralipid (10 μl/g BDW, l141-100mL, Sigma), and serum was collected at 0, 1.5, 3, and 6 hr for triglyceride, NEFA and glycerol assays. Glucose, insulin and triglyceride levels were measured using a Contour blood glucose monitor (9545C, Bayer) or an oxidase-peroxidase assay (Sigma P7119), insulin ELISA (Crystal Chem, Elk Grove village, IL, USA, #90080) and Infinity Triglycerides Reagent (Thermo Fisher Scientific TR22421). Glycerol and NEFA were measured by free glycerol reagent (F6428, Sigma) and free fatty acid quantification kits (Wako Diagnostics-NEFA-HR2), respectively. Blood lactate level was measured by lactate plus (Nova Biomedical). Pyruvate tolerance tests (PTTs) was performed by oral gavage of pyruvate (2.5 mg/g body weight) after overnight fasting. Serum FGF21, leptin, and adiponectin were determined by commercially available ELISA kits (for FGF21, cat# EZRMFGF21-26k, EMD Millipore; for leptin, cat# 90030, Crystal Chem; for adiponectin, cat# EZMADP-60K, EMD Millipore) following the manufacturers’ instructions, respectively.

### Mitochondrial respiration measurement

Adipose tissue oxygen consumption rate was determined using the Seahorse XFe24 Analyzer (Agilent) following the manufacturer-recommended BOFA (basal-oligomycin-FCCP-antimycin A/rotenone) protocol. The assay buffer was composed of 1 mM pyruvate, 2mM glutamine and 7mM glucose. Ex vivo mitochondrial function were measured by utilizing 6–8 mg adipose tissue. For tissues, oligomycin (10 μM), FCCP (8 μM) and antimycin A (4 μM) plus rotenone (2 μM) were added during the assay. OCR and extracellular acidification rate (ECAR) were recorded through the Seahorse instrument.

### Fatty acid measurements

For free fatty acid profiling analysis in serum (10 μL), sample preparation and derivatization was carried out as previously described ^28^. Total free fatty acid distribution in serum and tissue required alkaline hydrolysis prior to free fatty acid extraction and derivatization. Briefly, lipids from 5 μL of serum or an equivalent amount to 1 mg of tissue lipid extracts were extracted using the Folch method. The final organic extract (2 mL) was hydrolyzed by adding 150 μL of 1M KOH in MeOH followed by incubation at 55°C for two hours. The reaction mixture was then neutralized by the addition of 20 μL of acetic acid and dried down under nitrogen stream at 40°C. Samples were analyzed in a Nexera UHPLC LC-40 coupled to LCMS-9030 Q-TOF mass spectrometer (Shimadzu Scientific Instruments). Free fatty acid chromatographic separation was achieved on a Nexcol C18 1.8 μm 2.1×50 mm (Shimadzu Scientific Instruments) using a gradient elution with H_2_O 0.1% formic acid and MeOH:MeCN 1:1 (v:v) 0.1% formic acid.

### In vitro adipocyte differentiation

Stromal vascular fractions (SVFs) were isolated from mouse subcutaneous adipose tissue and cultured in DMEM/F12 medium containing glutamax (no. 10565-018, Gibco, ThermoFisher Scientific) supplemented with 10% FBS, penicillin–streptomycin and gentamicin (no. 15750060, Gibco, ThermoFisher Scientific) in a incubator with 10% CO_2_ at 37 °C. After SVF cells became confluent in the dish, the media was replaced with the differentiation initiating media that includes 0.5 mM 3-isobutyl-1-methylxanthine (IBMX), 1 μM dexamethasone, 5 μg/ml insulin and 1 μM rosiglitazone. After 48 h of incubation, SVF-derived adipocytes were maintained by the medium containing only 5 μg/ml insulin until day6 to day10.

### Proliferation assays

Cell proliferation was assessed through a crystal violet staining assay. Adipocytes (in vitro differentiated) were plated at a density of 10,000 cells per well, and collections were conducted every 2 days post-incubation. Subsequently, the harvested cells were washed with PBS, fixed for 10 minutes at room temperature using 10% formaldehyde, and stored in PBS at 4°C until all time points were gathered. The fixed cells were then stained with 0.1% crystal violet in a 20% methanol solution. After eliminating unincorporated stain through washing, crystal violet extraction was performed using 10% glacial acetic acid, and absorbance was measured at 595 nm.

### Liver cancer induction

Choline deficient diet with High fat (40 kcal%), high fructose (20 kcal%) and 2% cholesterol (RESEARCH DIETS, INC., D20030402) was provided at room temperature from 8 weeks old.

### LDH activity assay

Tissue LDH activity was measured by LDH Assay Kit (Abcam, ab102526, Cambridge, MA, USA) according to the manufacturer’s instructions. Briefly, 0.1 g of liver and 0.05 g of kidney tissues were homogenized in the LDH assay buffer followed by the centrifuge at 10,000 g for 15 min. Supernatants were diluted 1 :10,000 and assayed for LDH activity.

### NAD+ and NADH measurement

20mg of frozen kidney and liver tissue were homogenized in 400 μl of NADH/NAD Extraction Buffer. Tissue lysates were filtered through 10 kDa molecular weight cut off filters. NAD+ and NADH values were determined by NAD+/NADH Quantification Colorimetric Kit (BioVision, K337-100) according to the manufacturer’s protocol. 5 μl of tissue lysates were utilized for total NAD measurement instead of the original protocol. NAD^+^/NADH ratio was calculated by the value of total NAD^+^ and NADH (NAD^+^ = total NAD^+^ – NADH).

### Body composition analysis

The measurements of mouse whole-body compositions including total water, total fat mass and lean mass were performed by the EchoMRI whole body composition analyzer (Echo Medical Systems, Houston, Texas, USA).

### Intratracheal bleomycin injection

Anesthetize animals with isoflurane (2%) and create a small opening near the front part of the throat area. Dissect the platysma and anterior tracheal muscles to expose and reach the tracheal rings. Typically, administer intratracheal injection volumes at 2 μl per gram of weight. Load bleomycin (Sigma, B2434) solution into a 1mL subcutaneous syringe equipped with a 30-gauge, 5/16-inch needle. The dosage of the bleomycin is 1.5 unit/kg body weight. Maintain the syringe bevel side up and parallel to the trachea, injecting the entire instillate volume into the trachea. Withdraw the syringe from the trachea and apply surgical glue on the skin to close the opening.

### PPRE-luciferase assay

The luciferase reporter assay was performed using Bright-Glo Luciferase assay system (Promega, E2610). PPRE X3-TK-luc plasmid (addgene, #1015) was reverse transfected to the isolated SVF cells from subcutaneous adipose tissues with lipofectamine 3000 (Thermo Fisher Scientific, L3000001) before the induction of adipocyte differentiation. On day 6 of adipocyte differentiation, adipocytes were harvested for luciferase assay. The protein concentration of cell lysates was quantified by the BCA method (Thermo Fisher Scientific, A55865). Luciferase expression was normalized to protein concentration.

### Pyruvate uptake

Pyruvate [1-14C] (#NEC255050UC; PerkinElmer) was retro-orbitally injected (1 µCi per mouse) into mice after a 16 h fasting. Blood samples (0.15 ml) were then collected at 0, 10, 15 and 30 min after injection. At 30 min following injection, mice were euthanized, blood samples were taken and tissues were harvested. Tissues were quickly excised, weighed and frozen in liquid nitrogen and stored at −80°C until processing. Lipids were then extracted using a chloroform-to-methanol based extraction method ^29^. The radioactivity content of tissues, including blood samples, was quantified by scintillation counter (Tri-Carb 2910 TR, PerkinElmer, Waltham, MA).

### RNA-seq

RNA-sequence was performed by Novogene (Sacramento, CA, USA) by utilizing isolated RNA from the control kidney and adiponectin overexpressed kidney. After the QC procedures, mRNA from eukaryotic organisms is enriched from total RNA using oligo(dT) beads. For prokaryotic samples, rRNA is removed using a specialized kit that leaves the mRNA. The mRNA from either eukaryotic or prokaryotic sources then fragmented randomly in fragmentation buffer, followed by cDNA synthesis using random hexamers and reverse transcriptase. After first-strand synthesis, a custom second-strand synthesis buffer (Illumina) is added, with dNTPs, RNase H and Escherichia coli polymerase I to generate the second strand by nick-translation and AMPure XP beads is used to purify the cDNA. The final cDNA library is ready after a round of purification, terminal repair, A-tailing, ligation of sequencing adapters, size selection and PCR enrichment. Library concentration was first quantified using a Qubit 2.0 fluorometer (Life Technologies), and then diluted to 1ng/ml before checking insert size on an Agilent 2100 and quantifying to greater accuracy by quantitative PCR (Q-PCR) (library activity >2 nM). Libraries are fed into Novaseq6000 machines according to activity and expected data volume.

### Electron microscopy

Cells were fixed on prepared 25mm cover slips with 2.5% (v/v) glutaraldehyde in 0.1M sodium cacodylate buffer. After five rinses in 0.1 M sodium cacodylate buffer, they were post-fixed in 1% osmium tetroxide plus 1.5 % K33[Fe(CN)6] in 0.1 M sodium cacodylate buffer for 1 h in the dark at room temperature. Cells were rinsed with water and en bloc stained with 0.5% uranyl acetate in 25% MeOH overnight in refrigerator. After five rinses with water, samples were stained with 0.02M lead nitrate in 0.03M L-aspartate for 30 minutes in at 60 °C oven in the dark. Samples were dehydrated with increasing concentration of ethanol, infiltrated with Embed-812 resin and polymerized face up in a round histology mold in a 60 °C oven overnight. Hydrofluoric acid was used to remove glass from cover slip. Pieces of the epoxy disc were super glued to a blank block. Blocks were sectioned with a diamond knife (Diatome) on a Leica Ultracut UCT (7) ultramicrotome (Leica Microsystems) and collected onto copper grids, post stained with 2% Uranyl acetate in water and lead citrate. Images were acquired on a JEOL 1400+ transmission electron microscope equipped with a LaB6 source using a voltage of 120 kV and an AMT-BioSprint 16M CCD camera.

### Liquid chromatography/tandem mass spectrometry (LC/MS-MS)

LC/MS-MS was performed by UTSW proteomics core as described previously ^30^ to determine the proteins included in the subcutaneous adipose tissue lysate. Briefly, protein extraction was performed according to western blot section. Three biological replicates were processed and the average of each group was utilized to perform heatmap analysis. After protein separation by SDS page, gel samples were digested overnight with trypsin (Pierce) followed by reduction and alkylation with dithiothreitol and iodoacetamide (Sigma-Aldrich). After solid-phase extraction cleanup with Oasis HLB plates (Waters), the samples were analyzed by LC/MS-MS using an Orbitrap Fusion Lumos mass spectrometer (Thermo Electron) coupled to an Ultimate 3000 RSLC-Nano liquid chromatography system (Dionex). Samples were injected onto a 75 um i.d., 75-cm long EasySpray column (Thermo) and eluted with a gradient from 0 to 28% buffer B over 90 min. Buffer A contained 2% (v/v) ACN and 0.1% formic acid in water, and buffer B contained 80% (v/v) ACN, 10% (v/v) trifluoroethanol, and 0.1% formic acid in water. The mass spectrometer operated in positive ion mode with a source voltage of 1.5 kV and an ion transfer tube temperature of 275 °C. MS scans were acquired at 120,000 resolution in the Orbitrap and up to 10 MS/MS spectra were obtained in the ion trap for each full spectrum acquired using higher-energy collisional dissociation (HCD) for ions with charges 2-7. Dynamic exclusion was set for 25 s after an ion was selected for fragmentation. Raw MS data files were analyzed using Proteome Discoverer v2.4 SP1 (Thermo), with peptide identification performed using Sequest HT searching against the human-reviewed protein database from UniProt (downloaded April 8, 2022, 20361 entries). Fragment and precursor tolerances of 10 ppm and 0.6 Da were specified, and three missed cleavages were allowed. Carbamidomethylation of Cys was set as a fixed modification, with oxidation of Met set as a variable modification. The false-discovery rate (FDR) cutoff was 1% for all peptides.

### Bioinformatic analysis

Differential expression of genes between control and adiponectin KO mice were analyzed by use of genes having an fpkm of ≥0 in all samples. For the generation of heatmap, significantly changed protein coding genes were extracted from the original fpkm expression data. Hierarchical clustering was performed after normalization based on Z-score by Morpheus (https://software.broadinstitute.org/morpheus/). Scatter plot and volcano plot was generated by graphpad prism (Graphpad, San Diego, Calif., USA).

### PamGene Nuclear Hormone Receptor (NHR) Coregulator PamChip

The PPARγ coregulator binding profile was determined using the PamGene PamStation 12 (PamGene International) as previously described for PPARα ^31^. The tissue samples were pooled within their respective groups. They were lysed in Hinds’ NHR Coregulator Buffer supplemented with protease (Sigma) and phosphatase (Thermo Fisher Scientific) inhibitors using a TissueLyzer LT (Qiagen). The samples were centrifuged at 14,000 rpm for 10 minutes at 4°C, and the supernatant was transferred to a new tube. Protein concentration was determined using the Pierce BCA protein quantification kit (Thermo Fisher Scientific) and measured using a Varioskan Lux microplate reader (Thermo Fisher Scientific). Once the protein was extracted, 25 ng of protein was incubated with 25 nM PPARγ2 antibody (sc-166731) and 25 nM goat anti-mouse Alexa Fluor 488 (A-11001) secondary antibody while rotating at 10 RPM for 30 minutes at 4°C. The mixture was then added to a blocked NHR PamChip, and the PPARγ-coregulator binding was determined by taking fluorescent images over 102 cycles. The images were analyzed using the PamGene BioNavigator software. (PamGene International) and exported for data analysis. The intensity of the PPARγ-coregulator binding was normalized to the intensity of the reference point within each array.

### Statistical analysis

All data were expressed as mean ± SEM. Differences between two groups were examined for statistical significance by the Student’s t-test. Analysis of variance test (ANOVA) with a multiple comparison test using Prism software (Graphpad, San Diego, Calif., USA) was applied to the 4 group comparisons. *P* value <0.05 denoted the presence of a statistically significant difference. Each respective statistics methodology used is described in the respective Figure Legend.

## Results

### Overall phenotypes and adiponectin expression in adiponectin KO mice

The typical appearance of adiponectin KO pups during the lactation period is characterized by hair loss^32^. Hair loss can be observed from the rear to the neck, including the ventral and dorsal areas. Although this phenotype is common to all three types of adiponectin KO mice, the severity of hair loss varies depending on the specific line of adiponectin KO mice examined (**Fig. 1A**). To quantitate the severity of hair loss, we classified the severity of this phenotype into 3 groups: “*Normal*”, mild hair loss (“*Mild*”) and severe hair loss (“*Severe*”). *Normal* reflects no hair loss with an appearance resembling wild types in Fig. 1A. *Mild* represents less than 50% hair loss in the dorsal area, which can be observed in the Houston and Osaka adiponectin KO mice. *Severe* corresponds to more than 50% of hair loss, most typically seen in the Dallas adiponectin KO mice. 66% of Houston adiponectin KO mice show mild hair loss. None of the Houston mice show severe hair loss. On the other hand, the Osaka mice exhibit a mixed phenotype: 25.0% severe hair loss, 31% mild hair loss and 44% were normal (**Fig. 1B**).

**Figure 1.**
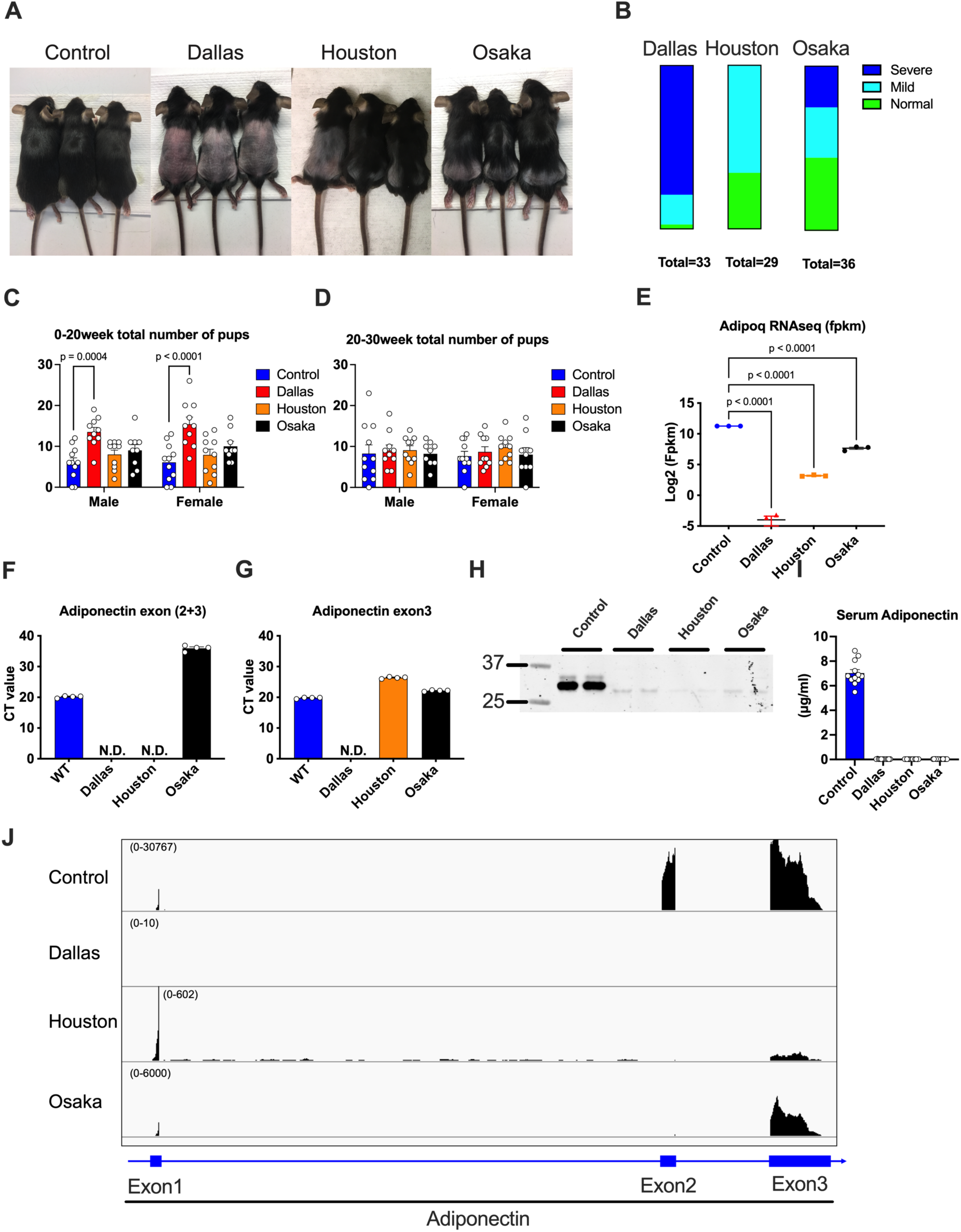
Overall phenotypes and adiponectin expression in adiponectin KO mice. **(A)** Representative appearance of wild type, Dallas, Houston and Osaka adiponectin KO mice at the age of 4 weeks. (n=3) **(B)** The severity of the hair loss of adiponectin KO mice was evaluated on the scale of normal, mild and severe. (n=29-36) **(C)** The number of pups in one breeding cage containing 1 male and 2 females from 0 to 20 weeks of age. (n=9-11) A two-way ANOVA was conducted followed by Dunnett’s multiple comparison test. **(D)** The number of pups in one breeding cage containing 1 male and 2 females from 20 to 30 weeks of age. (n=9-11) A two-way ANOVA was conducted followed by Dunnett’s multiple comparison test. **(E)** Adiponectin expressions in the subcutaneous adipose tissue based on the fpkm from RNA-seq (n=3) A one-way ANOVA was conducted followed by Dunnett’s multiple comparison test. **(F)** Quantification of adiponectin mRNA expression by primer pairs including exon2 and exon3. (n=4) **(G)** Quantification of adiponectin mRNA expression by exon3 primer pairs. (n=4) (**H**) Representative image of western blot of adiponectin in subcutaneous adipose tissues. (**I**) Serum adiponectin level in wild type and each adiponectin KO mouse (n=12-15) A one-way ANOVA was conducted followed by Dunnett’s multiple comparison test. (**J**) The representation of adiponectin expression in adiponectin locus in each adiponectin KO mouse by the Integrative Genomics Viewer (IGV) software. (n=3) Data are mean ±SEM. \**P*<0.05, \*\**P*<0.01, \*\*\**P*<0.001 and \*\*\*\**P*<0.0001

We counted the number of pups of dams at young age (0-20 weeks old) and older age of breeding (20-30 weeks old). At a young age, the Dallas adiponectin KO mice delivered more pups compared to the wild-types, as well as the Houston and the Osaka adiponectin KO mice (**Fig. 1C**). In contrast, we could not detect a significant difference in the number of pups born from dams from 20-30 weeks of age across all three knockout strains (**Fig. 1D**). At a young age, the Houston and Osaka adiponectin KO mice exhibit a slight (but not significant) increase of the number of pups. Overall, we conclude that adiponectin KO mice display a subtle increase in the number of pups, which is however not statistically significant. This increase is most pronounced in the Dallas adiponectin KO mice. To confirm that we are indeed dealing with knock-out mice, we also measured the expression level of adiponectin in subcutaneous adipose tissue by RNA-seq. Adiponectin KO mice showed lower levels of adiponectin compared to wild-type mice. Nevertheless, the Houston and Osaka adiponectin KO mice display residual expression of adiponectin, while the Dallas adiponectin KO mice completely lack any expression of adiponectin mRNA (**Fig. 1E**). To confirm this result, we performed quantitative PCR (qPCR) by utilizing two kinds of primers that detect exon 2 and exon 3 (exon 2+3) (**Fig. 1F**) and exon 3 only (**Fig. 1G**). Higher expression of adiponectin is detected in Houston and Osaka with the primer pairs specific for exon 3 compared to exon 2+3. This suggests that exon 3 is transcribed in the Houston and Osaka mice (**Fig. 1F and G**). Of note, the Osaka mice show 19 times higher expression of exon 3 than the Houston mice based on the CT value. However, in contrast to the adiponectin mRNA expression, we were unable detect any adiponectin protein in subcutaneous adipose tissue by Western blotting or by ELISA (**Fig. 1H and I**). This may reflect a relatively low-level expression of the protein, or there is a possibility that the anti-adiponectin antibodies used fail to detect the truncated adiponectin (**Fig. 1H**). We could also confirm the validity of the various adiponectin KO mouse models and the qPCR data by performing an alignment of RNA-seq results. In control mice, we could detect the peaks from exon 1, 2 and 3. The Dallas adiponectin KO mice do not show any peaks throughout exon 1 to exon 3. In contrast, consistent with the respective KO strategies, Houston and Osaka adiponectin KO mice lack only the exon 2 peak, but display the presence of mRNA species containing exon 1 and exon 3 (**Fig. 1J**).

### The Dallas adiponectin KO mice exhibit a lower body weight phenotype

To explore the metabolic phenotype under high-fat diet (HFD) conditions, we fed HFD starting at 8-10 weeks of age and performed the metabolic analysis after 2 months of HFD feeding. At the beginning of the HFD, no bodyweight differences were observed among adiponectin KO mice. However, Dallas adiponectin KO mice showed significantly lower body weight after 80 days of HFD treatment (**Fig. 2A and B**). Dallas KO mice showed no change in terms of lean mass. In contrast, the Houston KO mice exhibited the lowest lean mass (**Fig. 2C**). Consistent with the lower body weight, the Dallas KO mice showed a decreased fat mass, indicating that the body weight difference can be attributed to a reduction in fat mass (**Fig. 2D**). Specifically, the subcutaneous and brown adipose tissues exhibited lower weights, while the epididymal adipose tissue mass was unaltered in the Dallas KO mice (**Fig. 2E**). To explore the mechanism of the lower bodyweights and subcutaneous adipose tissue weights, we measured food intake, serum leptin and FGF21. However, these parameters were unchanged among the various adiponectin KO mice (**Fig. 2F, G and H**). Post 2 months of HFD, we performed systemic tolerance tests. For the oral glucose tolerance tests (OGTTs) and insulin tolerance tests (ITTs), the Dallas mice showed lower glucose and serum insulin levels, consistent with the leaner body weight phenotype (**Fig. 2I, J and K**). In contrast, the Osaka KO mice showed higher glucose and serum insulin levels during the OGTT and ITT (**Fig. 2I, J and K**). In pyruvate tolerance tests (PyTTs) which reflect the hepatic and renal gluconeogenic capacity, the Dallas KO mice show significantly lower blood glucose levels in response to pyruvate (**Fig. 2L**). This could be attributed to the lowered renal gluconeogenesis in Dallas adiponectin KO mice. Triglyceride tolerance tests (TGTTs) under HFD conditions exhibited higher serum TG levels in the Osaka and Houston KO mice, but not in the Dallas KO mice (**Fig. S1A**). On the other hand, TGTTs under insulinopenic conditions showed an elevation of serum TGs in the Dallas KO mice, but this phenomenon was not observed in the Osaka KO mice. This result suggests that the presence of even very small levels of adiponectin or one of its subfragments are vital for the retention of TG in adipocytes under insulinopenic conditions (**Fig. S1B**). This phenotype is not related to β adrenergic receptor activity, since a β adrenergic receptor agonist treatment, CL316,243 (CL), did not enhance lipolysis in the Dallas adiponectin KO mice (**Fig. S1C and S1D**). Consistent with the result of low retention of lipids in Dallas adipocytes, *in vitro* differentiation of adipocytes is hampered in the Dallas adiponectin KO mice (**Fig. S1E**). This is despite the fact that the cell proliferation is significantly higher in the stromal vascular fraction (SVF) in the Dallas KO mice (**Fig. S1F**). In line with the reduced fat pad weight in the Dallas KO mice, the average adipocyte size is significantly smaller (**Fig. 2M and 2N**). Lipid species composition in adiponectin KO subcutaneous adipose tissues reveals that Control and Houston KO mice share a comparable pattern of lipid species. In contrast, the Dallas adiponectin KO mice store overall less lipid species (**Fig. 2O and 2P**). Specific subtypes of lipid species, including adrenic acid and arachidonic acid, were enriched specifically in the Osaka mice (**Fig. 2O and 2Q**).

**Figure 2.**
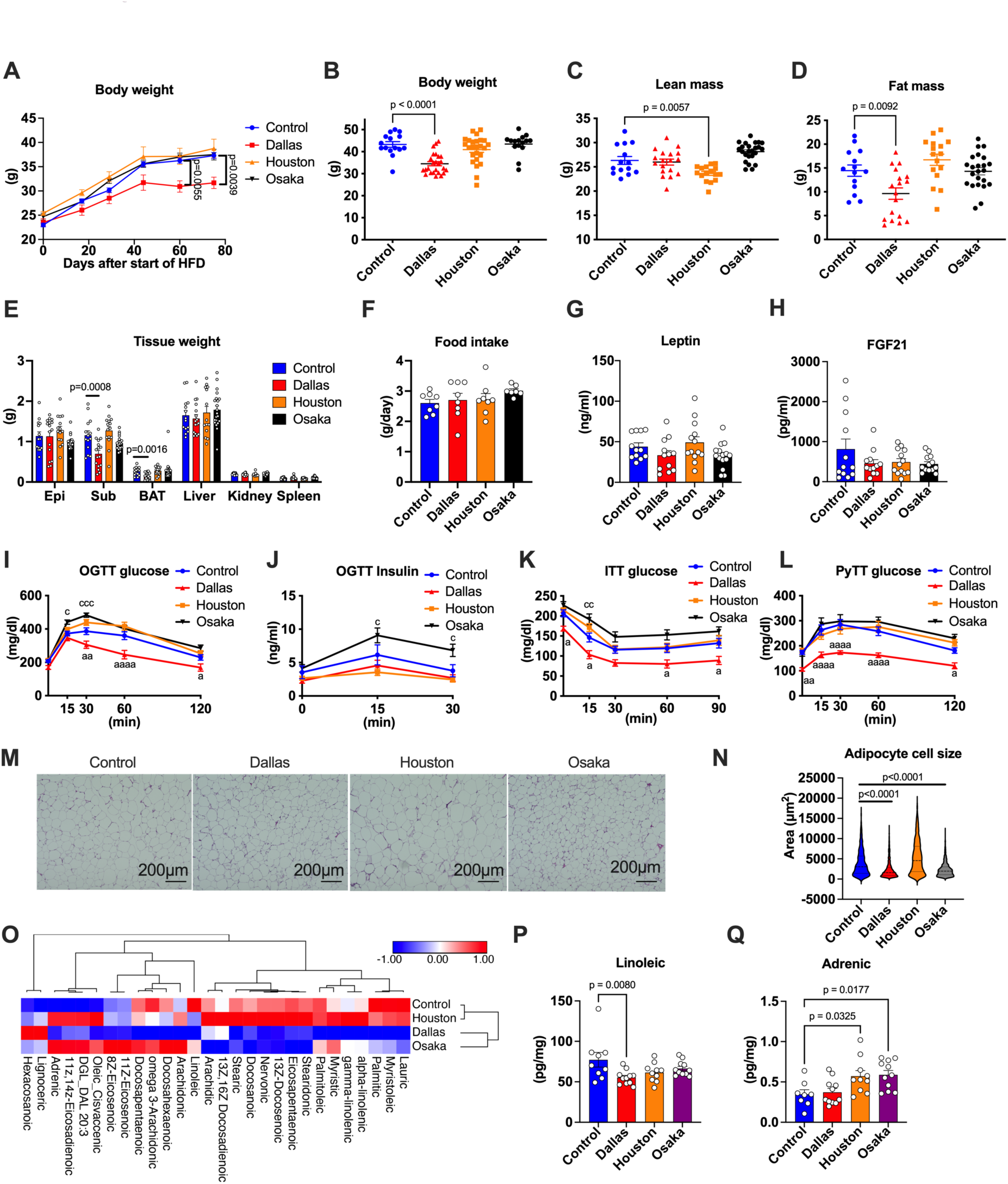
Dallas adiponectin KO mice exhibit lower body weight phenotype. Wild type and adiponectin KO mice were fed HFD from 8 weeks old. **(A)** Body weight of adiponectin KO mice after the start of HFD feeding (n=8-9). A two-way ANOVA was conducted followed by Dunnett’s multiple comparison test. **(B)** Body weight of adiponectin KO mice 3 months after the start of HFD feeding (n=15-25). A one-way ANOVA was conducted followed by Dunnett’s multiple comparison test. **(C)** Lean mass of adiponectin KO mice 3 months after the start of HFD feeding (n=15-25). A one-way ANOVA was conducted followed by Dunnett’s multiple comparison test. **(D)** Fat mass of adiponectin KO mice 3 months after the start of HFD feeding (n=15-25). A one-way ANOVA was conducted followed by Dunnett’s multiple comparison test. **(E)** Tissue weight of wild type and adiponectin KO mice including epididymal fat (Epi), subcutaneous fat (Sub), brown fat (BAT), liver, kidney and spleen. Tissues were harvested 4 months after the start of HFD. (n=14-21) A one-way ANOVA was conducted followed by Dunnett’s multiple comparison test. **(F)** Food intake of wild type and adiponectin KO mice. (n=7-8) **(G)** Serum leptin level in wild type and adiponectin KO mice 4 months after the start of HFD. (n=12-15) **(H)** Serum FGF21 level in wild type and adiponectin KO mice 4 months after the start of HFD. (n=12-13) (**I**) Blood glucose level in adiponectin KO mice during OGTT. (n=22-26) A two-way ANOVA was conducted followed by Dunnett’s multiple comparison test. **(J)** Serum insulin level in adiponectin KO mice during OGTT. (n=19-23) A two-way ANOVA was conducted followed by Dunnett’s multiple comparison test. **(K)** Blood glucose level in adiponectin KO mice during ITT. (n=17-22) A two-way ANOVA was conducted followed by Dunnett’s multiple comparison test. **(L)** Blood glucose level in adiponectin KO mice during pyruvate tolerance test. (n=24-29) A two-way ANOVA was conducted followed by Dunnett’s multiple comparison test. **(M)** Representative H&E staining images of subcutaneous adipose tissues in adiponectin KO mice. The scale bar indicates 200 μm. **(N)** Adipocyte size in adiponectin KO subcutaneous adipose tissue. (n=3) A Brown-Forsythe ANOVA was conducted followed by Dunnett’s T3 multiple comparison test. **(O)** Hierarchical clustering of the lipid species in adiponectin kO subcutaneous adipose tissues. (n=9-11) **(P)** Linoleic acid level in adiponectin KO subcutaneous adipose tissues. (n=9-11) A one-way ANOVA was conducted followed by Dunnett’s multiple comparison test. **(Q)** Adrenic acid level in adiponectin KO subcutaneous adipose tissues. (n=9-11) A one-way ANOVA was conducted followed by Dunnett’s multiple comparison test. \**P*<0.05, \*\**P*<0.01, \*\*\**P*<0.001 and \*\*\*\**P*<0.0001. Instead of asterisk, a, b and c is used to represent statistics for multiple comparisons. ^a^: Control versus Dallas, ^b^: Control versus Houston, ^c^: Control versus Osaka.

### Dallas KO mice exhibit higher mitochondrial function and the Osaka KO mice are characterized by higher fatty acid oxidation

The strong impact on subcutaneous adipose tissue in Dallas adiponectin KO mice prompted us to further investigate the adipose tissue phenotype in adiponectin KO mice. We performed RNA-seq of subcutaneous adipose tissue from each adiponectin KO flavor after 4 months of HFD feeding. We selected the 4440 protein coding genes whose False Discovery Rates (FDRs) are <0.05 in either of the comparisons between Control *vs* Dallas, Control *vs* Houston or Control *vs* Osaka. Additionally, according to the fold-change compared to the expression levels in the controls, we classified the genes into 3 categories, downregulated to less than 0.6 fold change (down), no change (nc), and upregulated by more than 1.4 fold change (up) groups. Then, we performed hierarchical clustering of these gene expressions in adiponectin KO mice. Hierarchical clustering revealed that the Control and Houston mice share similar gene expression signatures in their subcutaneous fat. In contrast, the Dallas and Osaka KO mice showed distinct gene expression patterns (**Fig. 3A**). Consistent with the hierarchical clustering, principal component analysis shows overlapping clusters for Control and Houston KO mice, whereas three segregated clusters are apparent amongst the Control, the Dallas and the Osaka transcriptomes (**Fig. 3B**). To explore the gene signatures in Dallas and Osaka subcutaneous adipose tissue, we performed the gene ontology analysis. The gene signatures enriched in Dallas mice are marked by elevated expression of mitochondrial respiratory chain components. This is also confirmed by the proteomics data (**Fig. 3C, D and S2A**). The Osaka gene signatures are typified by the presence of elevated levels of genes contributing towards fatty acid oxidation, even though these genes are also closely related to overall mitochondrial function (**Fig. 3C, E and S2B**). In line with these results, the subcutaneous adipose tissue oxygen consumption rates (OCRs) are higher in Dallas and Osaka mice when assessed in a Seahorse apparatus (**Fig. 3F and G**). Additionally, isolated mitochondria from Dallas and Osaka KO subcutaneous adipose tissues manifest significantly higher OCRs (**Fig. 3H and I**).

**Figure 3.**
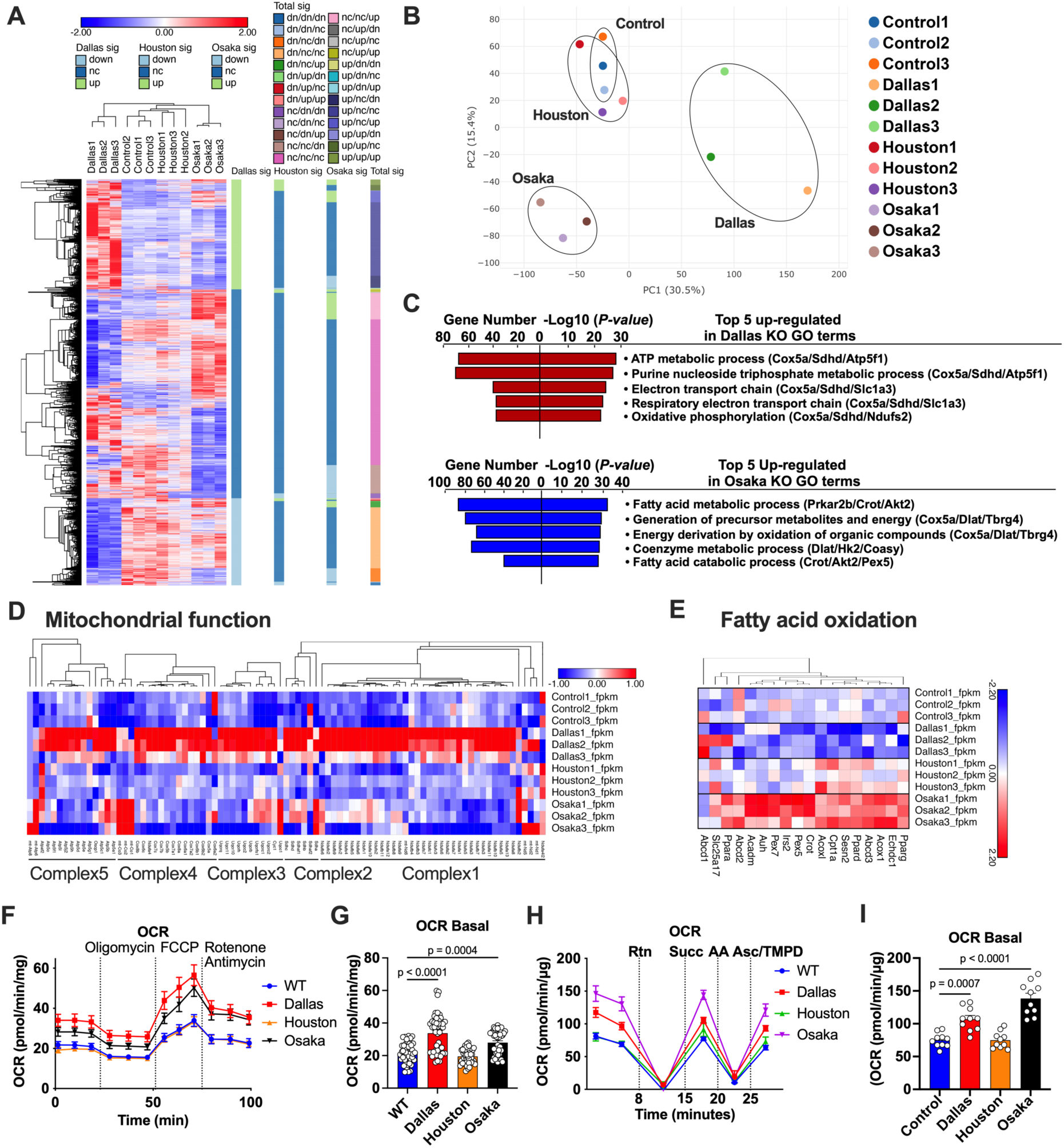
Dallas mice exhibit higher mitochondrial function and Osaka mice are characterized by higher fatty acid oxidation. Subcutaneous adipose tissues were harvested after 4 months of HFD feeding. 3 samples were pooled into one RNA-seq sample. 3 RNA-seq data in one group represent the transcriptome of 9 samples. **(A)** Hierarchical clustering of transcriptional profiles in control and adiponectin KO mice (n=3). **(B)** Principal component analysis of whole transcriptomes of adiponectin KO subcutaneous adipose tissues. (n=3) **(C)** Five most enriched pathway by gene ontology (GO) pathway analysis shows the characteristics of genes that were specifically up-regulated in Dallas (upper panel) and Osaka (lower panel) adiponectin KO subcutaneous adipose tissue. **(D)** Hierarchical clustering of the expressions of genes encoding mitochondrial components in adiponectin KO subcutaneous adipose tissues. (n=3) **(E)** Hierarchical clustering of the expressions of genes related to fatty acid oxidation in adiponectin KO subcutaneous adipose tissues. (n=3) **(F and G)** Oxygen consumption rate (OCR) of adiponectin KO subcutaneous adipose tissue at the Basal level followed by Oligomycin, FCCP and Rotenone/antimycin-A treatment (BOFA). The average values were obtained from 3 independent experiments. Each experiment contains 5 replicates in each group. A one-way ANOVA was conducted followed by Dunnett’s multiple comparison test. **(H and I)** OCR of mitochondria from each adiponectin KO subcutaneous adipose tissues at the Basal level followed by rotenone (Rtn), succinate (Succ), Antimycin-A (AA) and Ascorbate/TMPD treatment. A one-way ANOVA was conducted followed by Dunnett’s multiple comparison test. Data are mean ±SEM. \**P*<0.05, \*\**P*<0.01, \*\*\**P*<0.001 and \*\*\*\**P*<0.0001

### Adiponectin KO mice show lower serum lactate levels upon pyruvate gavage

We have shown that the different kinds of adiponectin KO mice exhibit significantly divergent phenotypes. This occurs despite the fact that circulating adiponectin is completely missing in all adiponectin KO mice. However, the non-secreted form of adiponectin that lacks signal sequence can exert effects intracellularly, particularly in the Houston and Osaka KO mice. Therefore, we were initially prompted to explore the phenotypes that the different KO strains had in common. This included a comparison of early-stage hair loss among these different null mice. This is presumably because the consistent phenotypes reflect the lack of circulating adiponectin rather than intracellular adiponectin. Along the same vein, we found that serum lactate levels were strongly suppressed in all adiponectin KO mice after a pyruvate gavage (**Fig. 4A**). To track the fate of the pyruvate, we gavaged ^14^C pyruvate, harvested tissues 15 min *post* gavage and determined the tissue radioactivity. The Dallas and Houston, but not the Osaka adiponectin KO mice, displayed higher levels of pyruvate uptake in the liver, heart, muscle and kidney (**Fig. 4B**). However, a higher incorporation of pyruvate to epididymal adipose was observed in the Osaka KO mice. Thus, pyruvate incorporation is higher in adiponectin KO mice across the board, even though there are distinct tissue preferences depending on the specific KO mouse that lead to the lower levels of circulating lactate (**Fig. 4B**). Since we observed the highest radioactivity in the kidneys per gram of tissue among the various organs, we focused more closely on the kidney pyruvate/lactate metabolism. The observation is that the Dallas and Houston adiponectin KO kidneys showed enhanced uptake of pyruvate associated with less lactate dehydrogenase (LDH) activity (**Fig. 4C**). This is the likely reason for the lower lactate production from pyruvate. Additionally, we observed significantly higher NADH levels in all adiponectin KO mice due to enhanced influx of pyruvate into the TCA cycle rather than being diverted to lactate (**Fig. 4D**). Among the urinary TCA cycle metabolites, citrate constitutes a major component associated with kidney stone formation^33^. Urine volumes along with urine citrate levels are significantly higher in the Dallas and Osaka adiponectin KO mice (**Fig. 4E and F**). Particularly, the Osaka mice display both disproportionately higher levels of urine volume and citrate, even if compared to the Dallas KO mice. This data suggests a prompt release of gavaged pyruvate as urinary citrate, which may explain the lower pyruvate uptake in the Osaka KO mice (**Fig. 4B**). In contrast to the lactate phenotype, lower urinary NH_4_ levels are more prominently seen in the Dallas adiponectin KO mice (**Fig. 4G**). Urinary NH_4_ levels are increased in the kidney tubular cell adiponectin overexpressing mice that we previously described (**Fig. 4H**). The readouts of higher citrate levels and lower NH_4_ levels in adiponectin KO mice along with the higher urinary NH_4_ level by renal adiponectin overexpression are very much consistent with the effects of pioglitazone, the well-characterized PPARγ agonist, to lower stone formation in patients ^34^, while PPARγ agonists are also amongst the best pharmacological stimulators of adiponectin production and release from adipocytes. Overall, lowered plasma lactate levels upon pyruvate gavage and higher NADH levels in the kidney are shared across all adiponectin KO mice and are caused by lowered LDH activity that results in an enhanced influx of pyruvate into the TCA cycle. In particular, the Osaka KO mice are characterized by a higher urine volume with high urine citrate level, resulting in a dramatic overall increased loss of citrate from the system. The hallmark of the Dallas KO mouse is its lower urine NH_4_. These subtle but physiologically very important differences highlight how profound the differential KO strategies affect the phenotypic characterization of the mice. This is due to the fact that we compare either the complete loss of adiponectin both in circulation and in the adipocyte vs. the loss of adiponectin in plasma with remnant adiponectin production within the intracellular milieu of the adipocyte (**Fig. 4I**).

**Figure 4.**
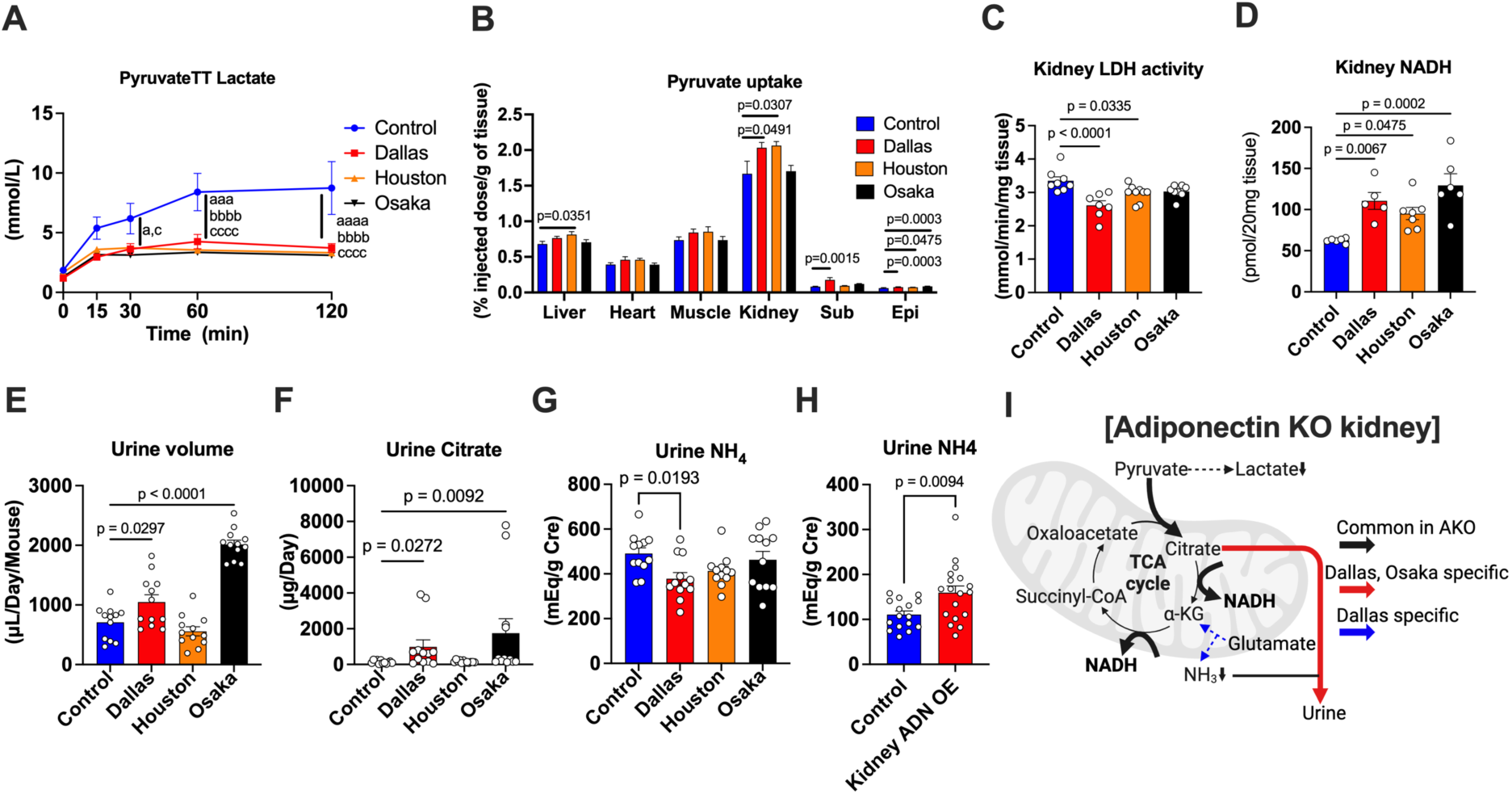
The phenotype that lowers serum lactate level upon pyruvate gavage is shared in all adiponectin KO mice. **(A)** Blood lactate level in adiponectin KO mice during pyruvate tolerance test. (n=8) A two-way ANOVA was conducted followed by Dunnett’s multiple comparison test. **(B)** The proportion of up-taken pyruvate by tissue out of total injected pyruvate. (n=7-8) A one-way ANOVA was conducted followed by Dunnett’s multiple comparison test. **(C)** LDH activity in the adiponectin KO kidney. (n=7-9) A one-way ANOVA was conducted followed by Dunnett’s multiple comparison test. **(D)** NADH level in the adiponectin KO kidney. (n=5-7) A one-way ANOVA was conducted followed by Dunnett’s multiple comparison test. **(E)** 24-hour urine volume in adiponectin KO mice. (n=12) A one-way ANOVA was conducted followed by Dunnett’s multiple comparison test. **(F)** 24-hour urine citrate level in the adiponectin KO mice. (n=12) A one-way ANOVA was conducted followed by Dunnett’s multiple comparison test. **(G)** Urine NH_4_ level in the adiponectin KO mice. (n=12) A one-way ANOVA was conducted followed by Dunnett’s multiple comparison test. **(H)** Urine NH_4_ level in the kidney tubular cell specific adiponectin overexpression mice (Kidney ADN OE). (n=12) An unpaired Student’s t-test was performed. **(I)** Schematic representation of the common and the distinct impacts of adiponectin deficiency on the kidney. Solid line and dash line represent the enhancement and down-regulation of the pathway, respectively. Data are mean ±SEM. \**P*<0.05, \*\*\**P*<0.001 and \*\*\*\**P*<0.0001. Instead of asterisk, a, b and c is used to represent statistics for multiple comparisons. ^a^: Control versus Dallas, ^b^: Control versus Houston, ^c^: Control versus Osaka.

### Dallas adiponectin KO mice are susceptible to DNA damage-mediated lung fibrosis

The anti-fibrotic properties of adiponectin are widely appreciated. However, it remains to be elucidated which tissues benefit the most from adiponectin, and whether there are differences in the loss of anti-fibrotic action amongst the different KO models. We used several types of stimuli to induce the fibrosis, such as intratracheal bleomycin in the lung (**Fig. 5**), HFD feeding (**Fig. S3**), kidney unilateral ureteral obstruction (UUO) (**Fig. S4**), and hepatic choline-deficient diet mediated hepatic cancer induction (**Fig. S5**).

**Figure 5.**
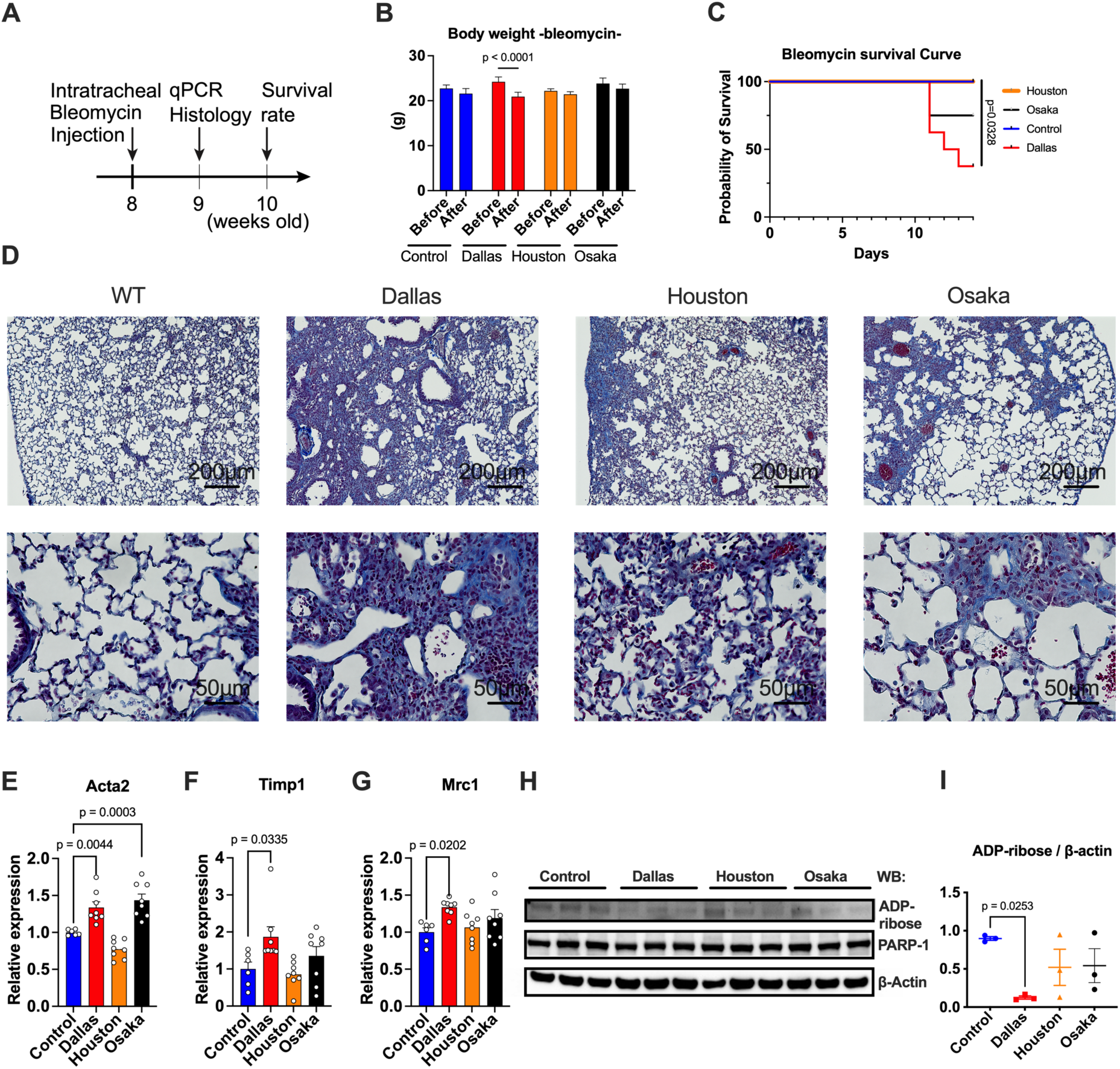
Dallas adiponectin KO mice are susceptible to DNA damage-mediated lung fibrosis. **(A)** Schematic representation of the time course of the intratracheal bleomycin injection. WT and adiponectin KO mice were subjected to the intratracheal bleomycin injection at the age of 8 weeks old. Lung tissues were harvested 1 week after the treatment for histology and qPCR. Survival rate was determined 2 weeks after the injection. **(B)** Body weight of adiponectin KO mice before and 1 week after the intratracheal bleomycin injection (n=6-8) A two-way ANOVA was conducted followed by Sidak’s multiple comparison test. **(C)** Survival curve of adiponectin KO mice after the treatment of bleomycin into the lung. (n=8) A Log-rank test with Bonferroni correction was performed. **(D)** Representative image of the trichrome staining of adiponectin KO lungs 1 week after the intratracheal bleomycin injection. The scale bar indicates 200 μm (upper panel) and 50 μm (lower panel), respectively. **(E)** Acta2 mRNA expression in the adiponectin KO lungs. (n=6-8) A one-way ANOVA was conducted followed by Dunnett’s multiple comparison test. **(F)** Timp1 mRNA expression in the adiponectin KO lungs. (n=6-8) A one-way ANOVA was conducted followed by Dunnett’s multiple comparison test. **(G)** Fn1 mRNA expression in the adiponectin KO lungs. (n=6-8) A one-way ANOVA was conducted followed by Dunnett’s multiple comparison test. **(H)** The image of the western blot of ADP-ribose and PARP-1 in adiponectin KO mice. (n=3) **(I)** The calculation of the density of western blot of ADP-ribose normalized by β-actin. (n=3) A one-way ANOVA was conducted followed by Dunnett’s multiple comparison test. Data are mean ±SEM.

We injected bleomycin to the lungs at 8 weeks of age followed by tissue harvest one week later for qPCR and histological analysis. Survival rates were calculated 2 weeks after the bleomycin injection (**Fig. 5A**). After one week post bleomycin injection, only Dallas adiponectin KO mice displayed a significant reduction in bodyweight (**Fig. 5B**). Subsequent to the weight loss, Dallas mice showed significantly lower survival rates. In contrast, the wildtype control mice and the Houston KO mice were totally protected from the bleomycin-mediated lung injuries (**Fig. 5C**). Trichrome staining revealed more fibrosis in the lungs of the Dallas KO mice. Compared to the Dallas KO mice, the fibrotic areas that occupy the lung alveoli are significantly reduced in both the Houston and Osaka KO mice (**Fig. 5D**). We assayed for the expression of fibrotic genes such as acta2, timp1 and fibronectin1 (fn1), all of which are significantly up-regulated in the Dallas and Osaka mice (**Fig. 5E, F and G**). Bleomycin induces single-strand DNA damage, which is accompanied by fibrosis development ^35,36^. Poly (ADP-ribose) polymerase1 (PARP1) is the enzyme that repairs single-strand DNA damage by inducing ADP-ribose ^37^. As one mechanism that might explain the enhanced fibrosis in the Dallas KO mice, these mice show a significantly reduced PARP1 activity after bleomycin injection, leading to decreased DNA repair in the Dallas mice, thereby causing more fibrosis upon bleomycin exposure (**Fig. 5H and I**). The conditions under which we observed the fibrotic phenotypes depend on the type of tissue examined, the experimental conditions used, and the type of KO mouse used. Under HFD conditions, Dallas KO mice display fibrotic phenotypes in adipose tissue, but not in the liver (**Fig. S3A and 3B**). While unilateral ureteral obstruction induces tubulointerstitial fibrosis compared to healthy control kidneys, we could not observe strong differences among the various adiponectin KO mice (**Fig. S4A and S4B**). To induce hepatocellular carcinoma, adiponectin KO mice were fed choline-deficient, high fructose, high cholesterol diets. One year after the start of this diet, overall, all adiponectin KO mice showed relatively higher body weights compared to WT mice (**Fig. S5A**). Even though the fat mass was not altered, the lean mass and the water content of the Dallas adiponectin KO mice was significantly higher than in the wild-type control mice (**Fig. S5B, C and D**). Consistent with this, the overall liver size and the liver tumor sizes were bigger in the Dallas adiponectin KO mice (**Fig. S5E, F and G**). Along with the hepatomegaly, the expressions of genes related to fibrosis were high in the Dallas KO mice, but low in the Osaka KO mice (**Fig. S5H**). Interestingly, gluconeogenesis-related genes were elevated in the livers of Osaka KO mice, which explains the deteriorated glucose metabolism observed in the Osaka KO mice (**Fig. S5I**). Picrosirius red staining indeed displayed higher fibrosis in the Dallas adiponectin KO mice compared to wild-type controls and Osaka KO mice (**Fig. S5J**).

### The intracellular overexpression of the globular form of adiponectin in the Dallas KO mouse partially recapitulates the Osaka adiponectin KO mice phenotypes

Full-length wild-type adiponectin comprises four domains: the signal sequence, a variable domain, a collagen domain, and a globular domain (**Fig. 6A Upper panel**). Since the Houston and Osaka KO mice lack exon 2 of adiponectin that encodes the signal sequence and the variable domain, we hypothesized that due to the presence of an alternate starting methionine in position 43, the globular form of adiponectin (gAPN) encoded in exon 3 remains intracellular and affects the cellular physiology of the adipocytes (**Fig. 6A**). The globular form of adiponectin forms a very stable trimer that we have previously crystallized and for which we solved the structure. We generated a transgenic construct starting with the second methionine of the adiponectin coding sequence that in position 43 (**Fig. 6A Lower panel**). To further explore the function of gAPN, we generated mice expressing this shortened form of adiponectin under the control of a tetracycline responsive element (TRE)*-gAPN*. The transcription of the gAPN mRNA is activated by breeding the mice into the background of an adiponectin rtTA mouse. This enables us to inducibly overexpress gAPN specifically in adipocytes in a doxycycline dependent manner. Considering the possibility that the effects of intracellular adiponectin can be saturated by the endogenous adiponectin expression in wild-type mice, we crossed the adiponectin-rtTA/TRE-gAPN mouse into the Dallas adiponectin KO background (gAPN) (**Fig. 6B**). Additionally, the gAPN coding sequence is modified to introduce a FLAG-tag at the C-terminus of the protein. This enables us to detect the gAPN protein with a FLAG-tag antibody. After one week of doxycycline treatment, we harvested the subcutaneous adipose tissue and extracted the proteins. We could successfully detect the globular adiponectin in the right size with a FLAG-tag antibody (**Fig. 6C**). In accordance with the notion that gAPN is lacking a signal sequence, we did not detect any globular adiponectin in the serum. Rather, we observed it exclusively in the subcutaneous tissue lysate (**Fig. 6D**). In order to define the metabolic phenotypes of gAPN, we fed HFD containing 600mg/kg doxycycline (HFDdox) for five months and then performed systemic metabolic tests (**Fig. 6E**). We started the HFDdox at the age of 9 weeks and monitored the body weights for the subsequent ten weeks. We could observe a higher body weight gain in gAPN than in the control mice (which are, in this case, the Dallas adiponectin KO mice) (**Fig. 6F**). Echo MRI showed that the body weight gain is in conjunction with the increased weight of fat mass (**Fig. 6G**). Specifically, the significant increase in fat mass was attributed to an increase in subcutaneous and brown adipose tissue rather than epididymal adipose tissue (**Fig. 6H**). gAPN mice exhibited higher glucose and serum insulin levels during an OGTT (**Fig. 6I and J**). gAPN mice showed higher glucose levels in an ITT, indicative of an insulin-resistant phenotype (**Fig. 6K**). Overall, the replenishment of gAPN into the Dallas adiponectin KO mice recovered the bodyweights and caused higher glucose levels. In contrast, gAPN showed a minimum impact on the lactate levels during a pyruvate tolerance test. While Adiponectin KO mice showed the expected lower lactate levels upon a pyruvate gavage (**Fig. 4A**), the reconstitution of adiponectin within adipocytes was insufficient to recapitulate the increase of lactate in circulation during the PTT (**Fig. 6L**). Principal component analysis (PCA) of RNA-seq revealed two distinct clusters that represent the distinct transcriptomes of the control mice (Adiponectin KO) and gAPN group (in the adiponectin KO background (**Fig. 6M**). gAPN overexpression suppressed browning of subcutaneous adipose tissues and enhanced fibrosis and inflammation, which may be responsible for the observed insulin resistance (**Fig. 6N, O and P**).

**Figure 6.**
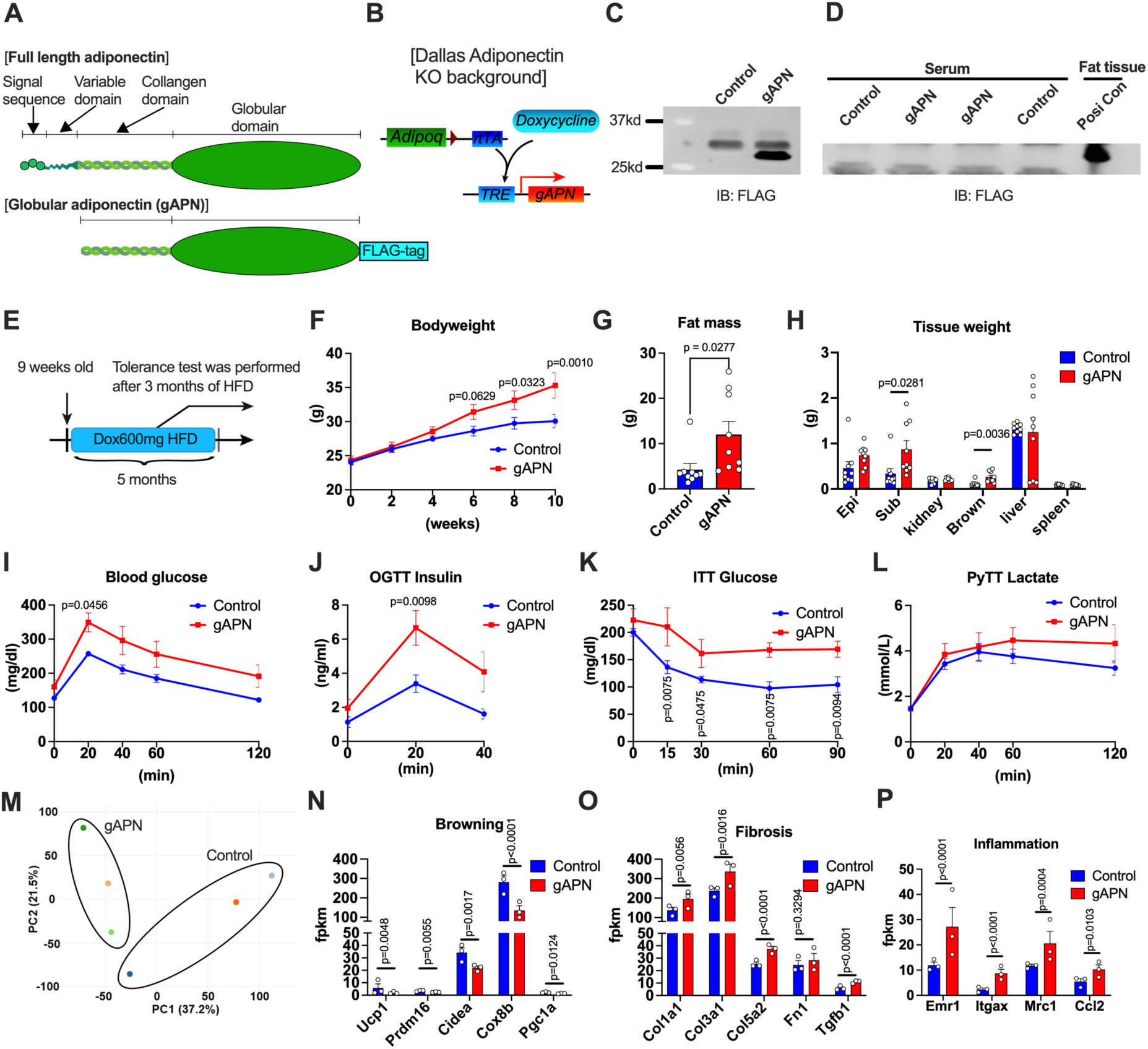
Intracellular form of adiponectin overexpression in Dallas partially recapitulates the Osaka adiponectin KO mice. **(A)** Full length adiponectin is composed of 4 domains including signal sequence, variable domain, collagen domain and globular domain. Overexpressed globular adiponectin lacks signal sequence and the former part of variable domain. FLAG tag is added to the C terminus of globular adiponectin. **(B)** By utilizing adiponectin rtTA and TRE-globular adiponectin transgene, globular adiponectin is overexpressed specifically in the adipocytes in the presence of doxycycline. Globular adiponectin is overexpressed under Dallas adiponectin KO background. **(C)** Globular adiponectin protein is determined in the adipose tissue by the FLAG tag antibody. **(D)** The serum level of globular adiponectin determined by western blot. **(E)** The schematic representation of the time frame of this study. Doxycycline 600mg/kg containing HFD was fed from 9 weeks old and kept for 5 months. Systemic tolerance tests were performed after 3 months of HFD feeding. **(F)** Body weight of control and globular adiponectin overexpression mice after the start of Doxycycline 600mg/kg containing HFD (control n=9, gAPN n=10). A two-way ANOVA was conducted followed by the two-stage linear step-up procedure of BKY correction. **(G)** Fat mass of control and globular adiponectin overexpression mice measured by Echo MRI (control n=9, gAPN n=9). An unpaired Student’s t-test was performed. **(H)** Tissue weight of each organ in control and globular adiponectin overexpression mice (control n=9, gAPN n=9). An unpaired Student’s t-test was performed. **(I)** Blood glucose level in globular adiponectin overexpression mice during OGTT (control n=9, gAPN n=10). A two-way ANOVA was conducted followed by Sidak’s multiple comparison test. **(J)** Blood insulin level in globular adiponectin overexpression mice during OGTT (control n=9, gAPN n=10). A two-way ANOVA was conducted followed by Sidak’s multiple comparison test. (**K**) Blood glucose level in globular adiponectin overexpression mice during ITT. (control n=7, gAPN n=4). A two-way ANOVA was conducted followed by the two-stage linear step-up procedure of BKY correction. (**L**) Blood glucose level in globular adiponectin overexpression mice during pyruvate tolerance test (control n=7, gAPN n=4). (**M**) RNA-seq was performed by utilizing subcutaneous adipose tissues after HFD doxycycline feeding. PCA analysis of the whole transcriptome of globular adiponectin overexpressed subcutaneous adipose tissues (control n=3, gAPN n=3). (**N**) The bar graph representation of the browning-related genes from RNA-seq data (control n=3, gAPN n=3). P-values were determined by use of DESeq2 R package. **(O)** The inflammation-related gene expression from RNA-seq data (control n=3, gAPN n=3). P-values were determined by use of DESeq2 R package. **(P)** The fibrosis-related gene expression from RNA-seq data (control n=3, gAPN n=3). P-values were determined by use of DESeq2 R package.

### Intracellular adiponectin regulates PPARγ transcriptional activity

To determine the intracellular distribution of gAPN, we performed immuno-electron microscopy (EM) utilizing differentiated adipocytes from SVF cells. The accumulation of immunogold particles represents the presence of gAPN, which is detected as the black dots. We observed a higher density of immunogold particles in the nucleus and in mitochondria compared to the control cells lacking gADN (**Fig. 7A**). In the nucleus, gAPN is more highly enriched in the heterochromatin area (**Fig. 7A right upper panel**). In mitochondria, gAPN is more enriched in the cristae structures (**Fig. 7A right lower panel**). To further define the role of gAPN localized to the nucleus, we immunoprecipitated gAPN using an anti-FLAG tag antibody and blotted with the PPARγ antibody. PPARγ protein is pulled down with globular adiponectin, specifically when the lysate is prepared from subcutaneous adipose tissue (**Fig. 7B**). This data suggest that globular adiponectin attaches to PPARγ, specifically in subcutaneous adipose tissue. To test the functional implications of this interaction of gADN with PPARγ, we determined PPARγ transcriptional activity with a luciferase construct driven by a PPAR response element (PPRE). Luciferase activity was up-regulated by gAPN, but only in the presence of rosiglitazone. This indicates that gAPN *per se* cannot drive PPARγ activity. Rather, gAPN activates PPARγ in concert with its agonist (**Fig. 7C**). Immunofluorescence staining of gAPN in subcutaneous adipose tissue further corroborates the fact that gAPN staining overlaps with mitochondria and – to a lesser extent – the nucleus (**Fig. 7D**). In agreement of a positive impact of PPARγ activity, in the Dallas KO, the degree of differentiation of adipocytes was significantly lower than in the wild-type controls (**Fig. 7E**) despite the fact that the PPARγ protein levels were comparable among wild-type and adiponectin KO adipocytes (**Fig. 7F**). To further explore the mechanisms of adiponectin to enhance PPARγ activity, we overexpressed globular adiponectin and PPARγ in HEK293 cells in the presence or in the absence of the PPARγ agonist rosiglitazone. We found that globular adiponectin is localized in the nucleus only in the presence of rosiglitazone (**Fig. 7G**). This is consistent with the activity assays shown in **Fig. 7C**. To verify the presence of gAPN in mitochondria, we performed immunoprecipitations of gAPN in lysates prepared from subcutaneous adipose tissue isolated from HFD fed mice by FLAG tag antibody, followed by proteomics. The top 20 proteins that were pulled down and more concentrated in gAPN subcutaneous adipose tissues compared to control included 15 types of mitochondrial proteins (**Fig. 7H**). Consistent with this observation, upon performing Seahorse activity assays on isolated mitochondria from gAPN subcutaneous adipose tissues, a significantly higher OCR was seen than in the wildtype controls (**Fig. 7I and J**).

**Figure 7.**
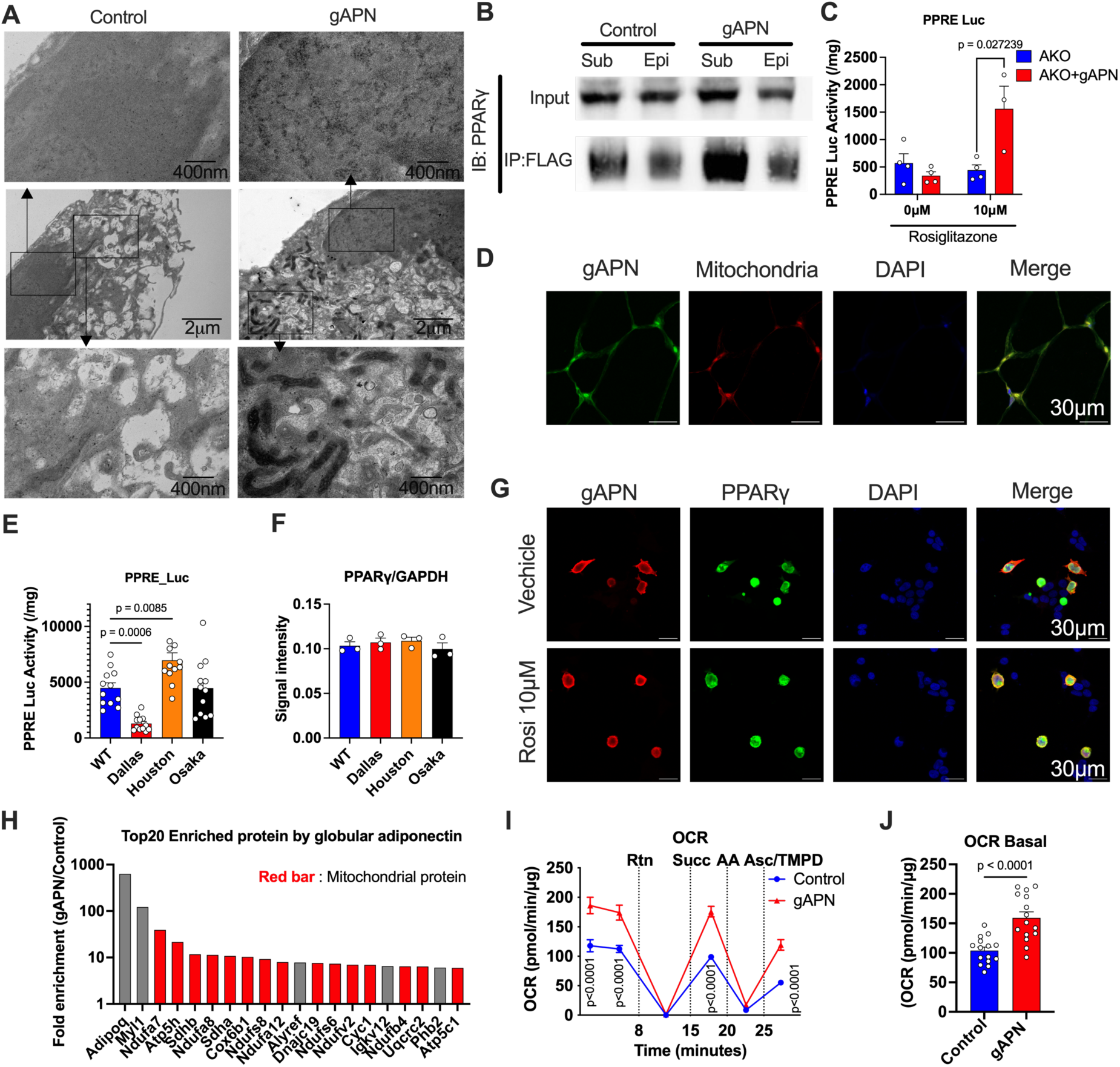
The distribution and functional involvement of intracellular globular adiponectin. **(A)** Electron microscopic view of differentiated adipocytes from SVFs from control (left panel) and gAPN (right panel). Overexpressed globular adiponectin with FLAG tag is stained by FLAG tag antibody and visualized by immunogold conjugated secondary antibody. The overview of adipocytes (middle panel), higher magnification of nucleus (upper panel) and mitochondria (lower panel). **(B)** Immunoprecipitation of epididymal and subcutaneous adipose tissue lysate from control and gAPN mice by FLAG tag antibody and blot by PPARγ antibody. The representative image of input and immunoprecipitated samples blot by PPARγ antibody. (n=1) **(C)** PPRE luciferase activity of the differentiated adipocytes with or without 10 μM rosiglitazone (n=3). An unpaired Student’s t-test was performed. **(D)** Immunofluorescence staining of globular adiponectin (gAPN) (green), mitochondria (red) and nucleus (blue) in the gAPN overexpressed subcutaneous adipose tissue. **(E)** PPRE luciferase activity of each adiponectin KO derived differentiated adipocytes in the presence of 10 μM rosiglitazone (n=12). A one-way ANOVA was conducted followed by Dunnett’s multiple comparison test. **(F)** The amount of PPARγ protein in the differentiated adipocytes derived from each adiponectin KO adipose tissue quantified by western blot (n=3). **(G)** Immunofluorescence staining of globular adiponectin (gAPN) (green), mitochondria (red) and nucleus (blue) with or without 10 μM of rosiglitazone in HEK293 cells. **(H)** Fold enrichment (gAPN/Control) of the proteins immunoprecipitated by FLAG tag antibody by use of subcutaneous adipose tissue lysate. **(I)** OCR of isolated mitochondria derived from control and gAPN subcutaneous adipose tissue at the Basal level followed by Rtn, Succ, AA and Asc/TMPD. A two-way ANOVA was conducted followed by Sidak’s multiple comparison test. **(J)** OCR of isolated mitochondria derived from control and gAPN subcutaneous adipose tissue at the Basal level. An unpaired Student’s t-test was performed. Data are mean ±SEM.

### Adiponectin modulates the recruitment of coregulators to PPARγ

To further investigate the mechanisms of the adiponectin-mediated regulation of PPARγ activity, we performed PamGene Nuclear Hormone Receptor (NHR) Coregulator Analysis of PPARγ. Wild type and Dallas AKO mice were treated with rosiglitazone for 2 weeks, followed by subcutaneous and epididymal adipose tissue harvests. The relative interactions of 155 coregulator proteins with PPARγ in wild-type and Dallas AKO subcutaneous fat pads induced by rosiglitazone (rosiglitazone-vehicle) are represented in a heatmap in alphabetical order (**Fig. 8A**). The bar graph shows the overall rosiglitazone-induced molecular signature of the coregulators interacting with PPARγ arranged in descending order from the left-hand side **(WT: Fig. 8B upper panel, Dallas: Fig. 8B lower panel)**. To compare the effect of adiponectin on the PPARγ coregulator binding profile, we quantified these graphs’ positive, negative, and total AUC. Rosiglitazone treatment positively recruited 70.2% out of the total AUC in WT mice, while 26.5% were positively recruited to the PPARγ coregulator complex in Dallas AKO mice. For the comparison between subcutaneous and epididymal adipose tissue, adiponectin deficiency had less of an effect on the PPARγ coregulator complex recruitment in epididymal adipose tissue **(Fig. S6A and B)**. In the presence of adiponectin, the binding of the Nuclear Factor kappa B (NFκB) Transcription Factor p65 subunit (TF65) with PPARγ is the most recruited coregulator to the PPARγ complex. At the same time, TF65 was the second most bound hit in the absence of adiponectin (**Fig. 8C**). Overall, the binding of coregulators to PPARγ is further enhanced by rosiglitazone in the presence of adiponectin. On the contrary, more proteins are disassociated from the PPARγ complex in the Dallas AKO subcutaneous adipose tissue by rosiglitazone treatment (**Fig. 8C**). A coregulator responsome plot revealed that TF65, NCOA1, PPRC, and PELP (*quadrant 4*) are regulated in opposite directions either with or without adiponectin. GNAQ and NCOA4 (*quadrant 3*) are down-regulated in the WT subcutaneous fat in response to rosiglitazone but further reduced in the Dallas AKO subcutaneous fat (**Fig. 8D**). Compared to subcutaneous adipose tissues, the epididymal adipose tissues show fewer alterations between WT and Dallas AKO mice. Notably, GNAQ is specifically down-regulated in Dallas AKO subcutaneous fat (**Fig. 8E**), yet up-regulated in the Dallas AKO epididymal fat. Instead, NCOA4 is the protein that is decreased mainly in the Dallas AKO epididymal fat **(Fig. S6E)**. These differentially regulated proteins within the two fat pads may explain why the subcutaneous fat tissue is more responsive to rosiglitazone than the epididymal fat with respect to PPARγ target gene expression (**Fig. S6C, D, and E**).

**Figure 8.**
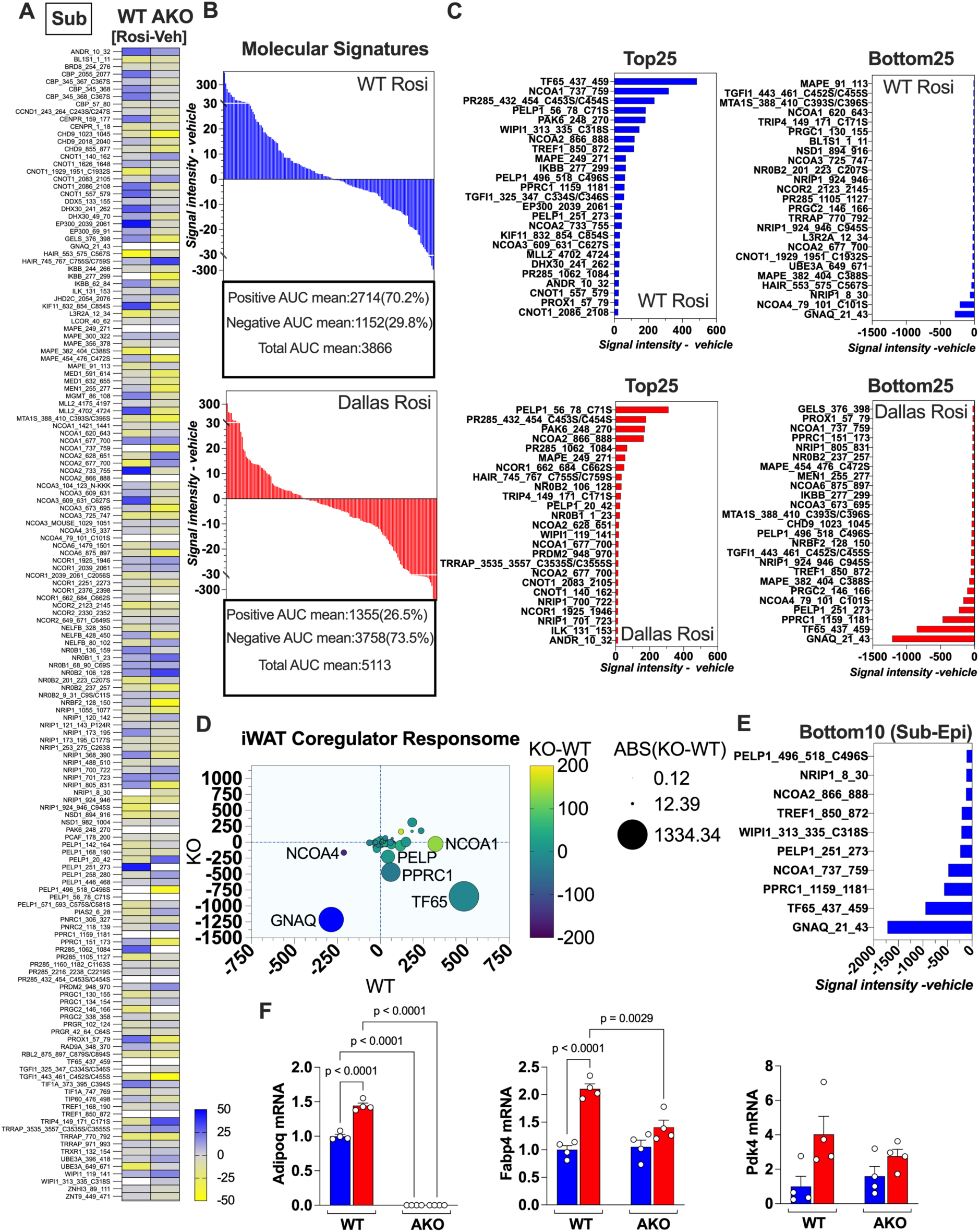
The loss of adiponectin alters the binding of coregulators to the PPARγ transcriptional complex in the subcutaneous adipose tissue. PamGene PamStation NHR analysis of the coregulators interacting with PPARγ in the subcutaneous adipose tissue (n=4 pooled). **(A)** Heatmap analysis showing the Rosiglitazone-induced (Rosi-vehicle) PPARγ coregulator interaction in the WT and Dallas AKO mice. **(B)** The Rosi-induced PPARγ molecular signature in the WT and Dallas AKO mice. **(C)** The top and bottom 25 coregulators are regulated by Rosi in each model. **(D)** The iWAT coregulator responsome compares the coregulator recruitment in the WT and AKO. **(E)** The bottom 10 coregulators interact with PPARγ in the AKO subcutaneous adipose tissue vs. epididymal adipose tissue. **(F)** Adipoq, *Fabp4*, and *Pdk4* mRNA expression in the WT and AKO mice treated with Rosi (n=4). A two-way ANOVA was conducted, followed by Tukey’s multiple comparison test. Data are mean ±SEM.

To confirm the effect of this alteration of the PPARγ coregulator complex on the transcription of genes, we measured the PPARγ target gene expressions such as adiponectin, *Fabp4*, and *Pdk4*. There was of course no mRNA expression of adiponectin in the subcutaneous and in the epididymal fat in the Dallas AKO mice. Fabp4 expression is significantly up-regulated by rosiglitazone in the WT subcutaneous fat, but not in the Dallas AKO fat. *Pdk4* exhibits the same tendencies but does not reach significance (**Fig. 8F**). In epididymal adipose tissues, the expression of these PPARγ target genes, *Fabp4* and *Pdk4*, are up-regulated to the same level by rosiglitazone in both WT and Dallas AKO **(Fig. S6F)**.

## Discussion

Over the years, we have noticed that distinct phenotypes have been reported for comparable experiments amongst the different adiponectin KO mice developed in different institutes by different cloning strategies. This prompted us to perform a head-to-head comparison and further explore the mechanisms underlying these discrepancies. By comparing the expression levels at the adiponectin locus by RNA-seq in each of the adiponectin KO mice, we observed a significant difference in the expression of exon 3 that encodes the bioactive part of adiponectin, the globular domain of adiponectin. In other words, the Dallas mice display a complete loss of adiponectin, while the Houston and the Osaka mice exhibited a lack of expression of exon 2 (leading to the elimination of *circulating* adiponectin as intended). However, they display residual expression of exon 3, in line with the KO strategy used for these mice ^12,14,15^, which leads to intracellular expression of globular adiponectin, a form of the protein that also likely exists in normal wildtype mice due to inefficient translocation into the ER of newly synthesized adiponectin. While these artifactual residual expression levels of the portion of the gene that remain intact are responsible for the differential phenotypes of the different KO mice, we learned valuable lessons in the process of analyzing this phenomenon and in fact believe that this globular domain is actually present in a number of cell types at low levels and is at least partially responsible for the phenotypic impact of adiponectin locally in a number of different cell types and tissues, as well as in the adipocyte proper as well.

The Dallas KO mice show a lower body weight and, associated with the lower body weight, improved glucose metabolism. However, the improved glucose metabolism is not only attributed to the lower body weight, because the Dallas mice show better insulin sensitivity even with comparable body weights to wild-type mice under normal chow diet conditions ^38^. One reason for this is that complete loss of adiponectin in the Dallas mice may impair renal gluconeogenesis ^39^, while the remnant expression of exon 3 in the Houston and Osaka mice enables them to maintain renal gluconeogenesis. This notion is also supported by the lowered urinary ammonium secretion in the Dallas adiponectin KO mice. Furthermore, all of the analysis here has been done on relatively young mice. At older ages, even the Dallas KO mice display metabolic dysfunction ^40^.

In terms of reproductive function, *in vitro* studies have shown the beneficial effects of adiponectin on fertility through controlling hormonal and reproductive systems ^41^. However, *in vivo* models, such as the adiponectin KO or AP2-driven ΔGly adiponectin overexpressing females, show only a minor impact on reproductive functions in PCOS models ^42^. In our study here, only the Dallas adiponectin KO mice displayed a higher number of pups. In part, this is also due to initiation of breeding earlier in life, while the reproductive abilities of the other adiponectin KO models are comparable to wild-type mice. These data suggest that remnant expression of intracellular adiponectin is sufficient to balance out the fertility phenotype of the Dallas adiponectin KO mice.

One of the obvious common phenotypes in adiponectin KO mice is hair loss. Indeed, adiponectin deficiency exacerbates age-related hair loss ^43^, but we also observe apparent hair loss in the neonatal period, a phenomenon which promptly recovers after weaning. This phenotype is caused by the composition of the maternal milk, containing a number of pro-inflammatory factors ^32^. This phenotype corresponds to the clinical observation that serum adiponectin level is lower in alopecia areata patients ^44^. It is notable that both adiponectin KO mice and ΔGly adiponectin overexpression mice exhibit neonatal hair loss^32^. This data corroborates the fact that an appropriate amount of adiponectin is vital for preventing neonatal hair loss. Since all of the adiponectin KO mice lack circulating adiponectin and manifest the hair loss phenotype, the hair loss phenotype in the adiponectin KO mice is predominantly dependent on circulating adiponectin. Nevertheless, the intracellular adiponectin in the Houston and Osaka adiponectin KO mice might be able to affect the inflammatory state of the mammary gland and, as a consequence, the severity of the neonatal hair loss, since the extent of the hair loss phenotype varies among adiponectin KO mice.

Obese adipose tissue is characterized by insulin resistance, chronic inflammation and fibrosis. Adiponectin prevents fibrosis systemically, including in the liver, kidney, skin, and even adipose tissue ^39,40,45,46^. Adiponectin also inhibits lung macrophage inflammation ^47^. A key transcriptional regulator for adiponectin is PPARγ. Agonists of PPARγ attenuate bleomycin-induced lung fibrosis ^48^ and also exert potent anti-fibrotic effects on the liver ^49^. Adiponectin prevents fibrosis through several mechanisms, such as preventing fibroblast activation to myofibroblasts ^50^ and inhibiting the nuclear factor kappa beta (NFκB) pathway ^51^. Here, we demonstrate that the Dallas KO mice are most susceptible to bleomycin-induced lung fibrosis. As one of the mechanisms responsible for the deteriorated lung fibrosis in the Dallas adiponectin KO mice is that polyADP-ribosylation is attenuated in the lungs of Dallas adiponectin KO mice. This data suggests that polyADP-ribose polymerase (PARP) activity is vital for single-strand DNA repair and is diminished in the Dallas adiponectin KO mice ^52^. Since bleomycin evokes both single- and double-strand DNA damage ^53^, loss of PARP activity worsens the bleomycin-mediated fibrosis. Since PARP acts intracellularly and in light of the fact that only Dallas adiponectin KO mice exhibited more fibrosis, this suggests that intracellular adiponectin may be crucial for maintaining NAD^+^ levels, PARP activity and proper maintenance of DNA repair.

Growing evidence demonstrates that lactate is a crucial metabolite for inter-organ cellular metabolism. The kidney and muscle are the primary sites for lactate production, even though cortical production of lactate can be offset by the consumption of lactate in the medulla ^54^. In line with this, pyruvate incorporation into organs is highest in the kidney per gram of tissue over the course of a pyruvate gavage. We have previously shown that renal tubular cell adiponectin overexpression increases the serum levels of lactate and other TCA cycle metabolites^39^. Consistent with this, adiponectin KO mice display lower circulating levels of lactate during a pyruvate tolerance test. Overall, the incorporation of pyruvate into organs is elevated in the adiponectin KO mouse. However, uptake into preferential organs varies amongst the different adiponectin KO mice. Additionally, lactate dehydrogenase (LDH) activity responsible for converting pyruvate into lactate is lowered across all adiponectin KO mice. These observations indicate more pyruvate influx into the TCA cycle at the expense of the conversion to lactate in adiponectin KO mice.

On the other hand, some of the kidney-related phenotypes are mouse line specific. The Osaka adiponectin KO mice are characterized by higher urine volume and higher citrate levels. This might also lead to a decrease of the serum lactate levels. Generally, urine NH_4_ levels reflect the use of amino acids and renal gluconeogenesis, a reaction that is specifically lowered in the Dallas adiponectin KO mice. This observation is further supported by the increase of NH_4_ in the kidney tubular cell specific adiponectin gain-of-function mice.

Adiponectin constitutes an important component of the fat-derived secretory proteome that exerts beneficial effects through membrane-bound receptors. However, a substantial amount of adiponectin is retained and distributed inside the adiponectin-producing cells ^55^. Intracellular adiponectin has been recognized to be retained at a higher level intracellularly in the context of metabolic challenges, such as HFD, tobacco smoke exposure or ovariectomies ^56–58^. However, whether intracellular adiponectin exerts distinct functional roles has not been investigated to date. Here, we report that exon 3 of the adiponectin gene encodes globular adiponectin. Upon removal of exon 2, as seen in the Houston and Osaka adiponectin KO mice, exon 3 is transcribed and translated. This portion of adiponectin lacks the signal sequence that is encoded in exon 2, thus preventing the secretion of adiponectin extracellularly. To explore the function of intracellular globular adiponectin in adipose tissue, we generated a doxycycline-inducible globular adiponectin construct and generated mice, specifically in adipocytes, and we did this in the background of the full Dallas KO. Overexpression of globular adiponectin in adipocytes increased the body weight and blood glucose levels during OGTTs and ITTs. Moreover, globular adiponectin overexpressing mice increased the oxygen consumption rate, very similar to the phenotype seen in the Osaka mice.

As a mechanism of a globular adiponectin function, immunoEM revealed that intracellular adiponectin mainly resides in mitochondria and in nuclei. Luciferase assays demonstrated that only the Dallas adiponectin KO mice (lacking the globular adiponectin portion as well) display lower PPARγ activity among the adiponectin KO mice (reflected in the smaller subcutaneous adipose tissue size in the Dallas adiponectin KO mice). On the contrary, globular adiponectin enhances PPARγ activity especially in the presence of the PPARγ agonist rosiglitazone, indicating an important contribution of globular adiponectin to PPARγ activity in adipocytes. Given that adipose tissue contains various endogenous PPARγ ligands, even in the absence of rosiglitazone, intracellular adiponectin should activate PPARγ and adipose tissue growth.

In terms of mitochondrial adiponectin, immunoprecipitation of the globular adiponectin highly enriches mitochondrial proteins and enhances mitochondrial oxygen consumption. Globular adiponectin itself seems to be a direct driver of mitochondrial respiration, but it could also be an indirect effect via PPARγ activation. The insulin resistant phenotype itself is not contradictory to the increase in mitochondrial respiration. We have previously generated mouse models with suppressed mitochondrial activity (induced by adipocyte mitoNEET overexpression), and these mice show higher insulin sensitivity ^59^.

To determine possible mechanisms of the reduced PPARγ activity in the Dallas AKO sub, we explored the rosiglitazone-induced PPARγ coregulator complex with or without adiponectin. We found that GNAQ and TF65 binding to PPARγ is strongly decreased due to an adiponectin deficiency in sub. The impact of adiponectin deficiency is much less apparent in epididymal fat, and GNAQ and TF65 are inversely regulated in epididymal fat compared to subcutaneous fat, and these two factors may have a key function in enhancing the adiponectin-mediated PPARγ activity. GNAQ encodes the G_q_ protein alpha subunit that couples to G protein-coupled receptors (GPCRs) to stimulate phospholipase Cβ (PLCβ) ^60^. Although the GPCRs involved in mediating the adiponectin-PPARγ-GNAQ complex interaction are unknown, the GNAQ-related signaling pathway could play an important role in PPARγ activation.

Importantly, given the much more pronounced effects of the loss of adiponectin in the subcutaneous fat compared to the epididymal fat, we may have found the mechanistic basis as to why PPARγ agonists in both preclinical as well as in clinical studies display a much stronger impact on the subcutaneous fat mass, leading to its preferential expansion – this is most likely due to the differential PPARγ interactome that adiponectin stabilizes in the subcutaneous adipocytes.

Taken together, we can explain the phenotypic discrepancies of the various adiponectin KO mouse lines via a head-to-head comparison in identical settings. We determined that some of the altered phenotypes derive from the residual expression of exon 3 in the Houston and Osaka adiponectin KO mice. We further suggest a mechanism of intracellular adiponectin that potentiates the PPARγ activity by modulating co-factors of PPARγ. This positive feedback loop of the adiponectin-PPARγ axis is fundamental to the differentiation and growth of adipocytes.

## Author contributions

T.O. designed and performed the experiments and acquired data with the help of Y.L., R.G., C.L. and X.N.S.. T.O. and P.E.S. interpreted the data and wrote the manuscript. C.M.K. helped the manuscript preparation. C.C. assisted western blot and immunoprecipitation. D.K. measured PARP activity. M.P. prepared histology and stained the slides. S.C. and M.Y.W performed intratracheal bleomycin injection. Z.A.K., E.A.B and T.D.H. determined the co-factors of PPARγ. J.S assisted luciferase assay. Q. Z., M.S. and R.K.G contributed to ChIP and IP experiment. Fatty acid in adipose tissue was measured by R.G.. I.S. and C.M.K, provided feedback on the manuscript. P.E.S. is the guarantor of this work and, as such, had full access to all the data in the study and takes responsibility for the integrity of the data and the accuracy of the analysis.

## Acknowledgments

We thank the UTSW Animal Resource Center, Metabolic Phenotyping Core, Transgenic Core, Live cell imaging core, Proteomics Core and Electron Microscopy Core. We also thank Shimadzu Scientific Instruments for the collaborative efforts in mass spectrometry technology resources. J.B.F. assisted making the constructs for NanoBit assay. This study was supported by US NIH grants RC2-DK118620, R01-DK55758, R01-DK099110, R01-DK127274 and R01-DK131537 to P.E.S and 1S10OD021685-01A1 to Electron Microscopy Core.

