## Supplemental information for "A comparison of adiponectin-deficient mice reveals the fundamental role of intracellular adiponectin"

### ***List of Supplement Materials***

1. Supplementary Figures S1 to S4

2. Tables S1 and 2

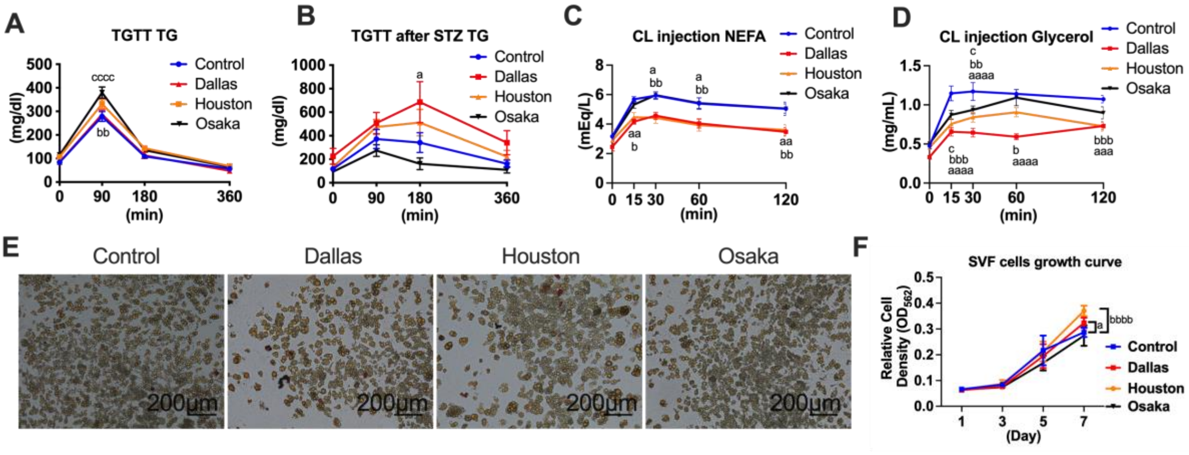


**Figure S1.** *Adipocyte size and profiles in adiponectin KO mouse lines*
**(A)** Blood TG level at different time point during TG clearance test. (n=23-30) A two-way ANOVA was conducted followed by Dunnett's multiple comparison test. **(B)** Blood TG level at different time point during TG clearance test after 1 week of STZ treatment. (n=7-13) A two-way ANOVA was conducted followed by Dunnett's multiple comparison test. **(C)** Blood NEFA level at different time point after CL compound injection. 0.5mg/kg CL316,243 was administered intraperitoneally after 2 months of HFD feeding and following overnight fasting. (n=8) A two-way ANOVA was conducted followed by Dunnett's multiple comparison test. **(D)** Blood glycerol level at different time point after CL compound injection. (n=8) A two-way ANOVA was conducted followed by Dunnett's multiple comparison test. **(E)** Representative image of Oil-Red O staining of in vitro differentiated adipocytes from adiponectin KO subcutaneous adipose tissues at the day 9 after the initiation of differentiation. Scale bar indicates 200 μm. **(F)** The time course of SVF cell proliferation. 8000 SVF cells from adiponectin KO subcutaneous adiposes were seeded on the culture dish. (n=6) A two-way ANOVA was conducted followed by Dunnett's multiple comparison test.

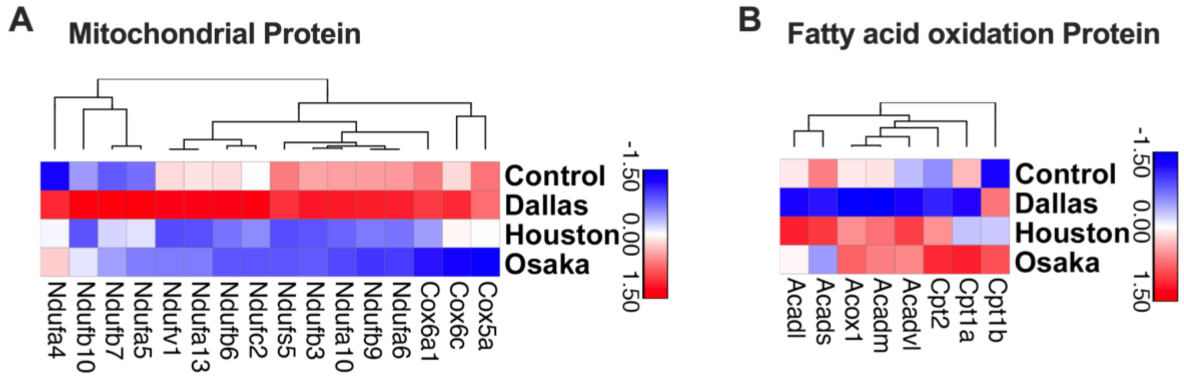


**Figure S2.** *Proteome analysis of mitochondrial functions*
**(A)** Hierarchical clustering of the mitochondrial protein. Subcutaneous adipose tissues from adiponectin KO mice after 4 months of HFD were utilized for proteomics. (n=3) **(B)** Hierarchical clustering of the fatty acid oxidation proteins. (n=3)

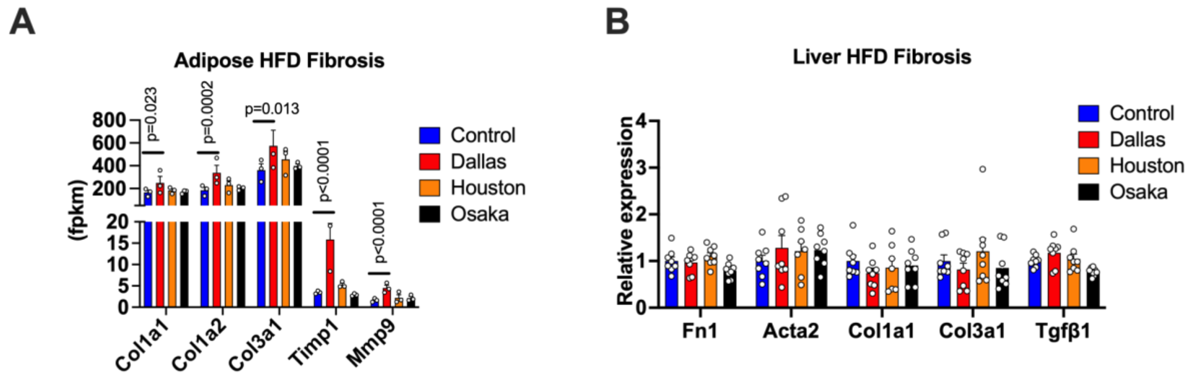


**Figure S3.** *The impact of adiponectin deficiency on adipose and hepatic fibrosis under HFD condition*

Subcutaneous adipose tissues and livers from adiponectin KO mice were harvested after 4 months of HFD feeding. **(A)** Fibrosis related gene expression in the adiponectin KO subcutaneous adipose tissues from RNA-seq data. (n=3) P-values were determined by DESeq2 R package followed by Bonferroni correction. **(B)** Fibrosis related gene mRNA expression in the adiponectin KO liver. (n=7-8)

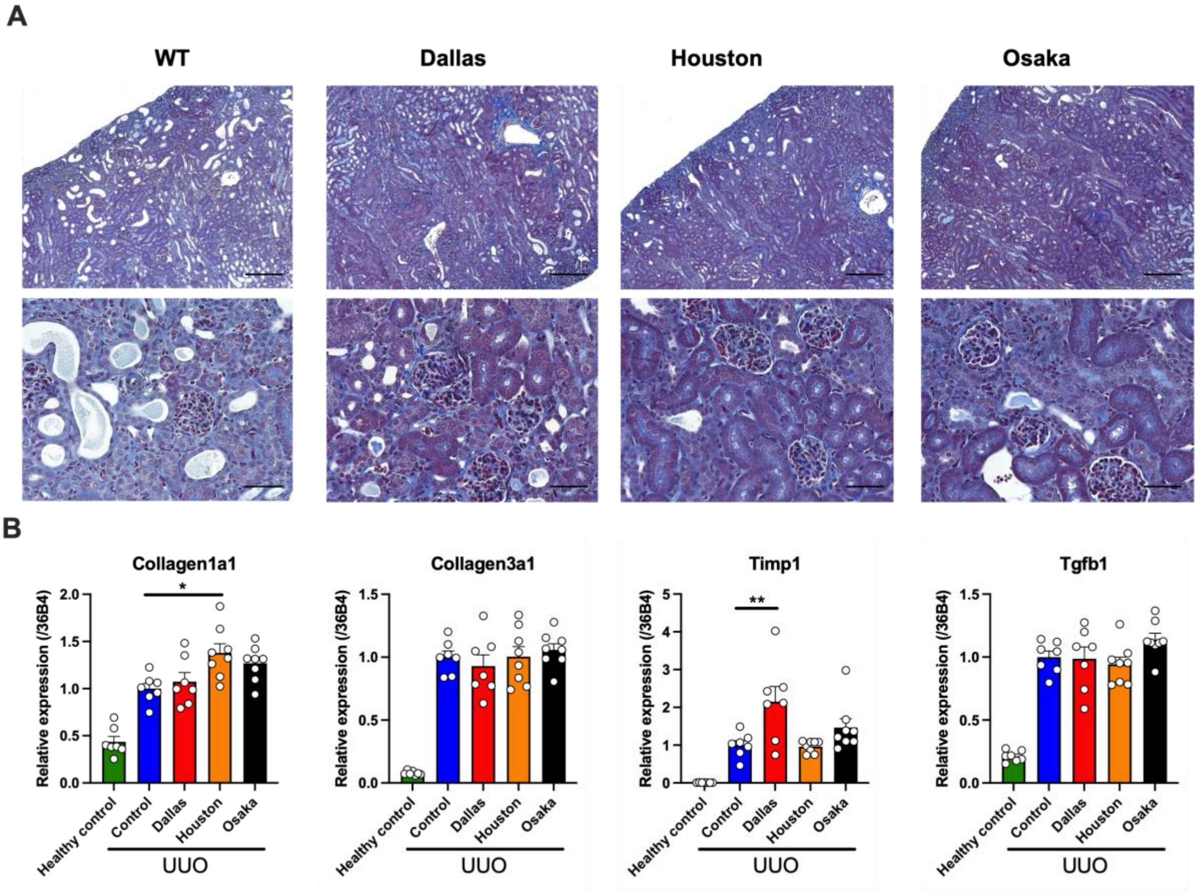


**Figure S4.** *The impact of adiponectin deficiency on renal fibrosis by unilateral ureter obstruction (UUO)*

The kidney of adiponectin KO mice were harvested 1 week after UUO. **(A)** Representative trichrome staining image of adiponectin KO kidney tissues after UUO. Scale bars of upper and lower panel indicate 200 and 50 μm, respectively. **(B)** Expression of genes involved in fibrosis including Collagen1a1, Collagen3a1, Timp1 and Tgfβ1. A one-way ANOVA was conducted followed by Dunnett's multiple comparison test.

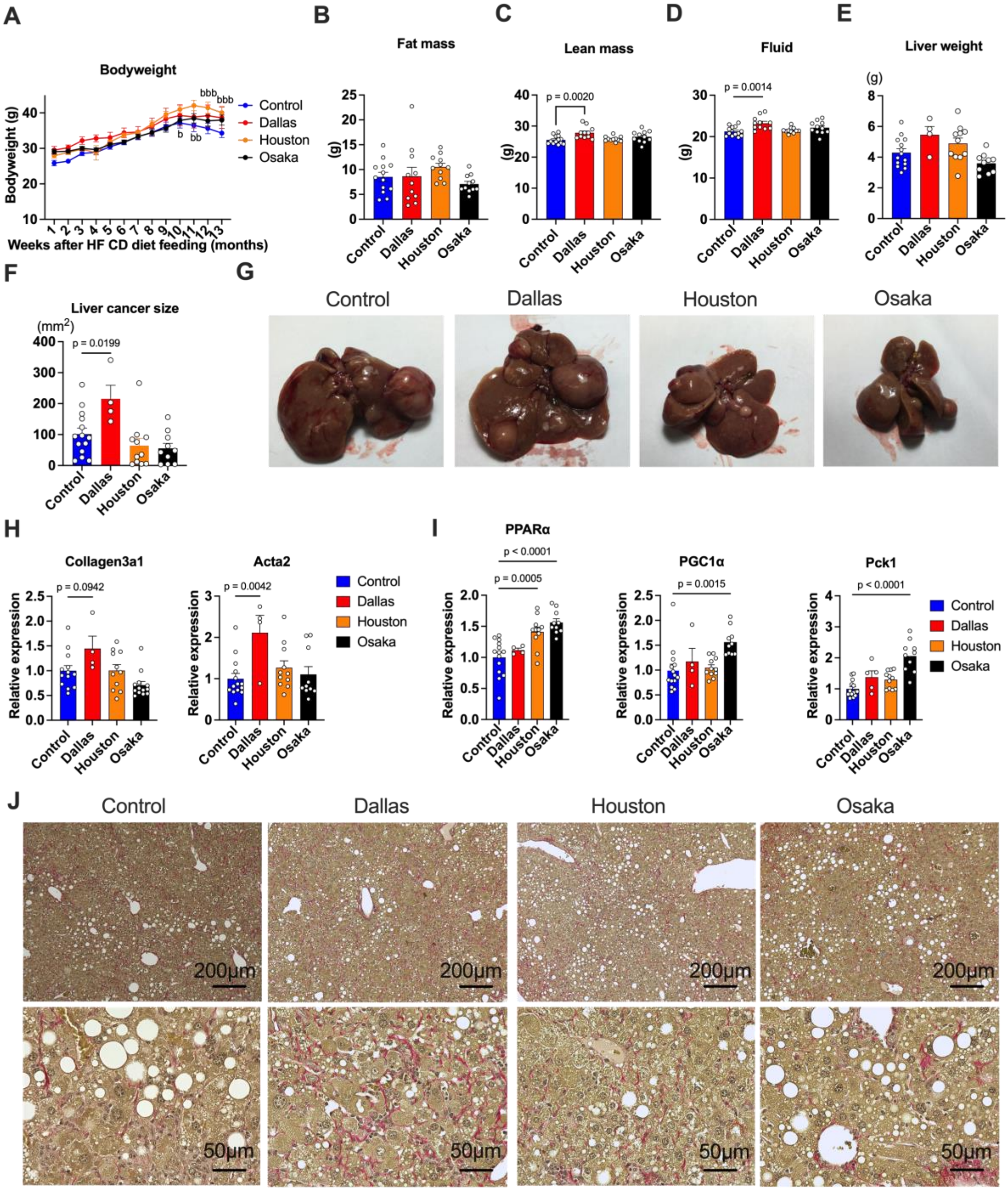


**Figure S5.** *Dallas adiponectin KO mice are prone to develop liver cancer by choline deficient high fructose high cholesterol HFD*

**(A)** Body weight of adiponectin KO mice under the treatment of choline deficient high fructose high cholesterol HFD. (n=6-14) **(B)** Fat mass of adiponectin KO mice. (n=11-13) **(C)** Lean mass of adiponectin KO mice. (n=11-13) A one-way ANOVA was conducted followed by Dunnett's multiple comparison test. **(D)** Fluid weight of adiponectin KO mice. (n=11-13) A one-way ANOVA was conducted followed by Dunnett's multiple comparison test. **(E)** Liver weight of adiponectin KO mice. (n=4-14) **(F)** Liver cancer size of adiponectin KO mice. Total liver cancer size in each mouse was calculated by summing up the size of multiple cancers. (n=4-14) A one-way ANOVA was conducted followed by Dunnett's multiple comparison test. **(G)** The representative image of the whole liver in adiponectin KO mice. **(H)** The expressions of genes involved in fibrosis including Collagen1a1, Collagen3a1, Tgfβ1, Timp1 and Acta2. (n=4-14) A one-way ANOVA was conducted followed by Dunnett's multiple comparison test. **(I)** The expressions of genes involved in gluconeogenesis including PPARα, PGC1α, Pck1. (n=4-14) A one-way ANOVA was conducted followed by Dunnett's multiple comparison test. **(J)** Representative image of picrosirius red staining of the liver in adiponectin KO mice.

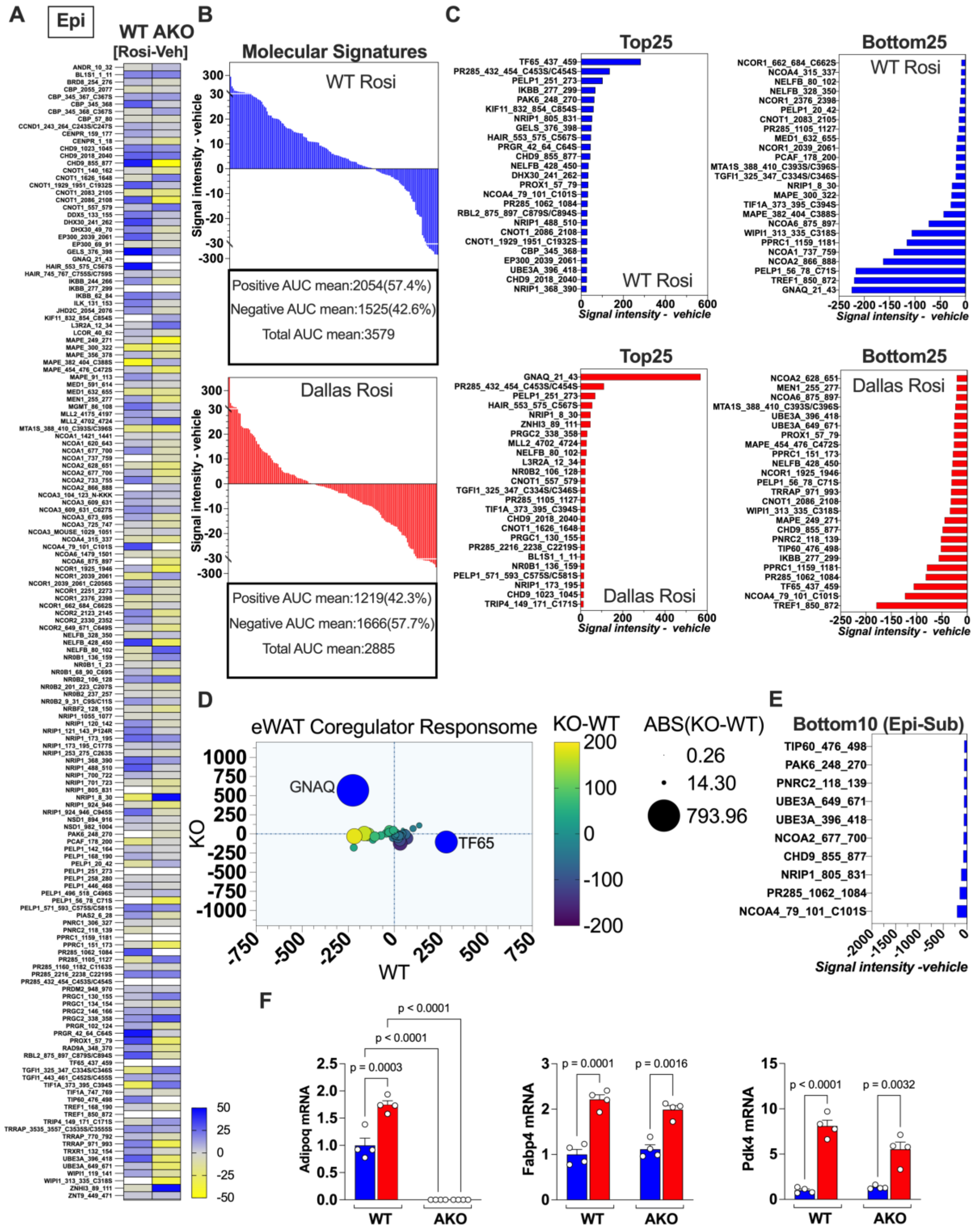


**Figure S6.** *The loss of adiponectin alters the coregulators to the PPARγ transcriptional complex but not affects the target gene expression in the epididymal adipose tissue.*

PamGene PamStation NHR analysis of the coregulators interacting with PPARγ in the epididymal adipose tissue (n=4 pooled). **(A)** Heatmap analysis showing the

Rosiglitazone-induced (Rosi-vehicle) PPARγ coregulator interaction in the WT and Dallas AKO mice. **(B)** The Rosi-induced PPARγ molecular signature in the WT and Dallas AKO mice. **(C)** The top and bottom 25 coregulators are regulated by Rosi in each model. **(D)** The iWAT coregulator responsome compares the coregulator recruitment in the WT and AKO. **(E)** The bottom 10 coregulators interact with PPARγ in the AKO subcutaneous adipose tissue vs. epididymal adipose tissue. **(F)** Adipoq, Fabp4, and Pdk4 mRNA expression in the WT and AKO mice treated with Rosi (n=4). A two-way ANOVA was conducted followed by Tukey's multiple comparison test. Data are mean ±SEM.

Table S1

### List of genotyping primer sequences

| Mouse strain | Forward primer | Reverse primer |
|---|---|---|
| Adiponectin-rtTA | TGCAGGTCCTGATTGGATGTG | TTTCCTTGTCGTCAGGCCTTC |
| TRE-Adiponectin | CTCCTGGAGAGAAGGGAGAGAAAG | CCGTGATGTGGTAAGAGAAGTAGTAGAG |
| Dallas Adiponectin KO WT | TTGGACCCCTGAACTTGCTTCACACC | TCCTGAGTTCAATTCCCAGCACCCAC |
| Dallas Adiponectin KO KO | TTGGACCCCTGAACTTGCTTCACACC | GGATGCGGTGGGCTCTATGGCTTC |
| Houston Adiponectin KO WT | GGTGGCTCACAACCATTCA | CTCCCAGGAGGTCTTCATCA |
| Houston Adiponectin KO KO | GGTGGCTCACAACCATTCA | CTTCCTGACTAGGGGAGGAG |
| Osaka Adiponectin KO WT | ATGAAGACCTCCTGGGAGAGTG | AGGAGCTAGCTCTTCAGTTG |
| Osaka Adiponectin KO KO | AAGAACTCGTCAAGAAGGCGATAGAAGGCG | AGGATCTCCTGTCATCTCACCTTGCTCCTG |

### Table S2

### List of qPCR primer sequences

| Gene name | Forward primer | Reverse primer |
|---|---|---|
| Acta2 | GTACCACCATGTACCCAGGC | GCTGGAAGGTAGACAGCGAA |
| Timp1 | CCCCAGAAATCAACGAGACCA | ACTCTTCACTGCGGTTCTGG |
| Fn1 | GCCCTGGTTTGTACCTGCTA | GGAATCTTTAGGGCGCTCAT |
| Col1a1 | GCTCCTCTTAGGGGCCACT | CCACGTCTCACCATTGGGG |
| Col3a1 | CTGGAGAACCTGGTGCAAAT | CCTCGGAAGCCACTAGGAC |
| Tgfβ1 | ACCATGCCAACTTCTGTCTG | CGGGTTGTGTTGGTTGTAGA |
| Emr1 | TTACGATGGAATTCTCCTTGTATATCAT | CACAGCAGGAAGGTGGCTATG |
| Itgax | CTGAGAGCCCAGACGAAGACA | TGAGCTGCCCACGATAAGAG |
| Mrc1 | TGTGGTGAGCTGAAAGGTGA | CAGGTGTGGGCTCAGGTAGT |
| Ccl2 | CCACTCACCTGCTGCTACTCAT | TGGTGATCCTCTTGTAGCTCTCC |
| PPARα | GCCTGTCTGTCGGGATGT | GGCTTCGTGGATTCTCTTG |
| Pgc1a | CAGCCTCTTTGCCCAGATCT | CCGCTAGCAAGTTTGCCTCA |
| Pck1 | CCACAGCTGCTGCAGAACA | GAAGGGTCGCATGGCAAA |
